# centroFlye: Assembling Centromeres with Long Error-Prone Reads

**DOI:** 10.1101/772103

**Authors:** Andrey V. Bzikadze, Pavel A. Pevzner

## Abstract

Although variations in centromeres have been linked to cancer and infertility, centromeres still represent the “dark matter of the human genome” and remain an enigma for both biomedical and evolutionary studies. Since centromeres have withstood all previous attempts to develop an automated tool for their assembly and since their assembly using short reads is viewed as intractable, recent efforts attempted to manually assemble centromeres using long error-prone reads. We describe the centroFlye algorithm for centromere assembly using long error-prone reads, apply it for assembling the human X centromere, and use the constructed assembly to gain insights into centromere evolution. Our analysis reveals putative breakpoints in the previous manual reconstruction of the human X centromere and opens a possibility to automatically close the remaining multi-megabase gaps in the reference human genome.

---

Long-read technologies (such as Pacific Biosciences and Oxford Nanopore) have greatly increased the contiguity of genome assemblies as compared to short-read technologies. However, although the existing long-read assemblers, such as Falcon (Chin et al., 2016), Miniasm (Li, 2016), Flye (Kolmogorov et al., 2019), HINGE (Kamath et al., 2017), Canu (Koren et al., 2017), Marvel (Nowoshilow et al., 2018), and wtdbg2 (Ruan and Li, 2019), have been used in many sequencing projects, they typically fail to resolve long segmental duplication (Vollger et al., 2019) and long tandem repeats (Kolmogorov et al., 2019). This paper focuses on the latter challenge – assembling long tandem repeats, and specifically on centromere assembly, the problem that was viewed as intractable until recently. As shown in Kolmogorov et al., 2019, the existing long-reads assemblers are inaccurate even in the case of relatively short *bridged tandem repeats* that are spanned by long reads. Although Flye improved on other tools in assembling such repeats (Kolmogorov et al., 2019), assembly of *unbridged tandem repeats* remains an open problem.

Centromeres are among the longest tandem repeats in the human genome and the biggest gaps in the reference human genome assembly. As a result, studies of associations between sequence variations and genetic diseases currently ignore ≈3% of the human genome. This is unfortunate since centromeres play critical roles in chromosome segregation and a large component of genetic disease result from aneuploidies arising during meiosis (Nagaoka et al., 2012). In addition, variations in centromeres are linked to cancer and infertility (Enukashvily et al., 2007, Sahin et al., 2008, Ting et al., 2011, Ferreira et al., 2015, Giunta and Funabiki, 2017, Black et al., 2018, Smurova and De Wulf, 2018, Barra and Fachinetti, 2018, Zhu et al., 2018, Miga 2019). Centromere sequencing is also important for addressing open problems about centromere evolution (Schueler et al., 2001, Alkan et al., 2007, Shepelev et al., 2009, Lower et al., 2018). These studies revealed extremely fast evolutionary development of centromere: complex higher-order centromeric repeats are unique to the hominid lineage and are missing even in such close species as gibbons (Cellamare et al., 2009). Recent discovery of large archaic blocks of Neanderthal DNA spanning human centromeres (*cenhaps*) reveals the potential of centromeres for studies of human population history (Langley et al., 2019).

Human centromeres represent long tandem repeats (also known as *satellite DNA*) that are often repeated thousands of times with extensive variations in copy numbers in the human population. Although long read technologies facilitated analysis of some centromeres (Miga et al., 2014, Jain et al., 2018), no software tool for centromere reconstruction has been released yet and it remains unclear how accurate the centromere reconstructions that resulted from previous semi-manual efforts are. To close all remaining gaps in the human genome (including centromeres) the recently established Telomere-to-Telomere consortium aims to generate its first complete assembly (Miga et al., 2019).

Below we describe the centroFlye algorithm for centromere assembly and the automatic reconstruction of the centromere on chromosome X (referred to as cenX). We provide evidence that the previous manual cenX reconstruction have missed a large fraction of cenX and resulted in errors that were fixed in our new assembly. Analysis of this assembly revealed that human X chromosome is partitioned into repeat subfamilies and provided initial insights into centromere evolution. Although this paper is limited to reconstructing a single cenX centromere, our next goal is to extend centroFlye for reconstructing all human centromeres.

## Results

### Centromere architecture

The *alpha satellite* repeat family forms ≈3% of the human genome (Hayden et al., 2013). Each alpha satellite is formed by *monomers*, each approximately 171 nucleotides in length. Consecutive monomers form *higher-order repeat* (*HOR*) *units* in which blocks of multiple monomers form a larger domain that can be repeated thousands of times. Individual monomers within a HOR show low similarity to each other (50–90% sequence identity) while HORs within a single centromere show high sequence similarity (95%-100% sequence identity). Organization and nucleotide sequence of HORs is specific for a particular chromosome. Figure 1 shows a dot-plot of the consensus HOR sequence on the X centromere (referred to as DXZ1) formed by 12-monomers with length 2,055. HORs typically occupy multi megabase-sized segments that include rearrangements and transposon insertions (Sevim et al., 2016).

**Figure 1.**
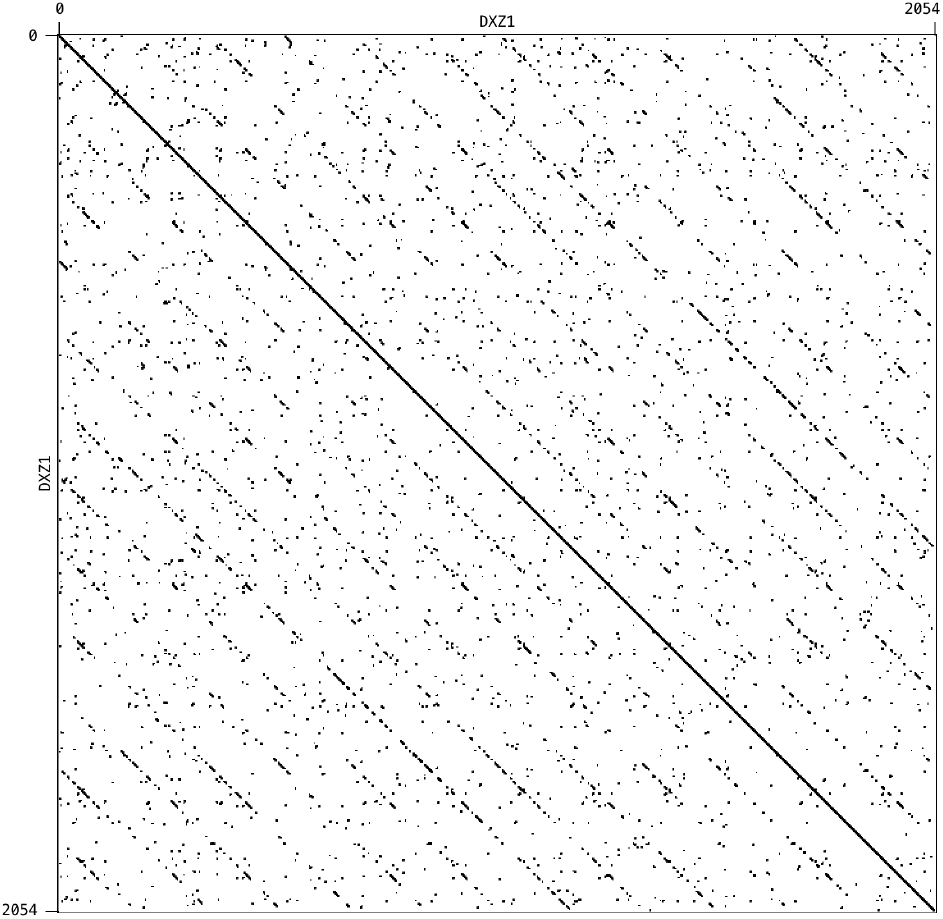
A dot-plot of a DXZ1 reveals twelve monomers. (GenBank: X02418.1). The sequence identity between different monomers varies from 60% to 86%. The dot-plot was constructed using the Gepard tool (Krumsiek et al., 2007) with the “word length” = 6.

Since centromere assembly of a diploid genome is particularly challenging, studies of the centromeres on X and Y chromosomes in the male genome represent a somewhat simpler (albeit still very complex) problem (Mahtani and Willard, 1998, Schueler et al., 2001, Schindelhauer and Schwarz, 2002). Miga et al., 2014 constructed a set of cenX-specific reads and used it to model the monomer organization of cenX. This model enabled analysis of variant sites on cenX, information that can be used for extending cenX model to cenX assembly.

### centroFlye pipeline

centroFlye takes a set of long error-prone reads from the entire genome and a consensus HOR (characterizing a specific centromere) as an input. It further classifies a read as *centromeric* if it contains at least one HOR and assembles the centromeric reads into a centromere sequence.

centroFlye modifies the approach to resolving *unbridged repeats* used in the Flye assembler (Kolmogorov et al., 2019) for the case of tandem repeats. Flye finds *divergent positions* within an unbridged repeat (positions where repeats copies differ from each other) and uses them as stepping stones for resolving these repeats. However, since the mapping of reads within cenX is unknown, it is unclear how to infer the divergent positions between various copies of a HOR. centroFlye instead defines a set of *rare k*-mers (*k*-mers that appear in a single of a few copies of a HOR) and uses them for reconstructing centromeres.

The centroFlye pipeline consists of the following steps (Figure 2):

**Figure 2.**
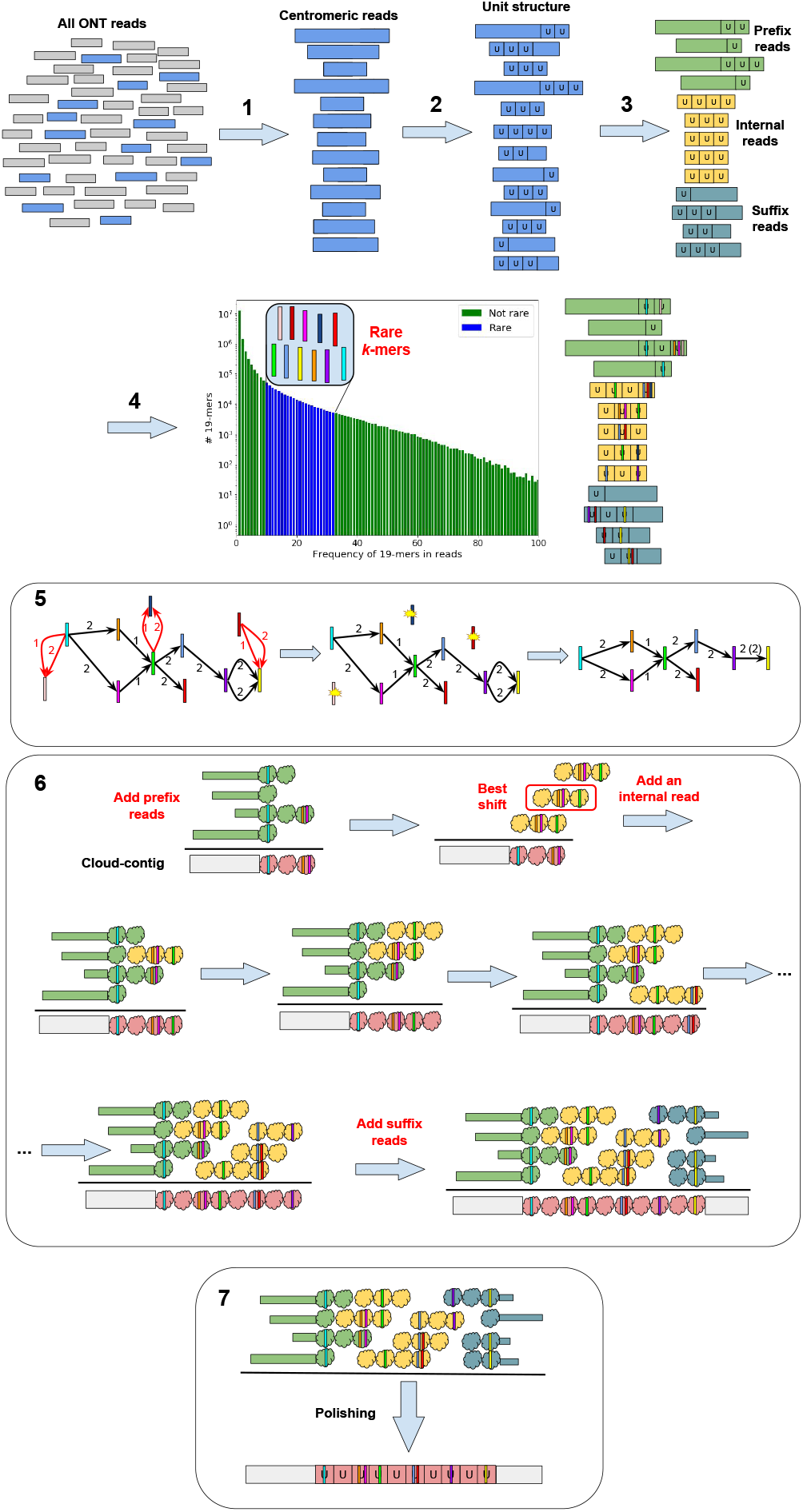
centroFlye pipeline. **1:** Recruitment of centromeric reads from the entire read-set. **2:** Partitioning each read into units, where each unit represents a HOR copy. **3:** Classifying centromeric reads into prefix, internal, and suffix reads. **4:** The frequency histogram of all *k*-mers in centromeric reads reveals rare *k*-mers. Each colored vertical bar represents a rare *k*-mer. **5:** Construction of the distance graph. Rare *k*-mers represent vertices and each edge connects a pair of rare *k*-mers occurring in the same read. Edge labels represent distances (in units) between rare *k*-mers in a read. An edge is red if there are conflicting parallel edges connecting corresponding vertices, and black otherwise. (Left) The distance graph constructed on all rare *k*-mers. (Middle) The distance graph with conflicting parallel edges and isolated vertices removed. (Right) The final distance graph with collapsed multi-edges. **6:** Reconstruction of a centromere. Each unit in each read is represented as a cloud with colored bars representing unique *k*-mers. After all prefix reads are added to a cloud-contig, the best alignment (shift) of each read against the cloud-contig is selected and the read with the highest-scoring alignment is added to the growing cloud-contig. This procedure is repeated until the suffix reads are added to the cloud contig. **7:** Polishing the reconstructed centromere sequence.

1. recruiting centromeric reads,
2. partitioning centromeric reads into *units*, where each unit represents a HOR copy,
3. classifying centromeric reads into *prefix* reads (that start before the centromere and “enter” it), *internal* reads (that are contained entirely within the centromere), and *suffix* reads (that start in the centromere and “leave” it),
4. identifying *rare centromeric k-mers*,
5. constructing the *distance graph* to filter out false positives among rare centromeric *k*-mers,
6. reconstructing the centromere,
7. polishing the reconstructed centromere sequence.

In Methods we describe each of these steps and illustrate them using the cenX assembly.

### Dataset

We analyzed the dataset of Oxford Nanopore reads generated by the Telomere-to-Telomere consortium (Miga et al., 2019) and released on March 2, 2019. The dataset contains reads generated from the CHM13hTERT female haploid cell line. There are 11,069,717 reads (155 Gb total length, 50x coverage) with the N50 read length equal to 70 kb.

7,651,424 out of 11,069,717 reads with length below 5 kb (1.7x coverage of the human genome) have limited impact on the centromere assembly. In contrast, 999,562 *ultralong* reads (longer than 50 kb) have the biggest impact on the centromere assembly and result in ≈32x coverage of the human genome.

### centroFlye and Telomere-to-Telomere assemblies of cenX

We analyzed the centroFlye assembly, the Telomere-to-Telomere consortium assembly v0.4 released on March 2, 2019 (referred to as *T2T4 assembly*), and its latest version (v0.6) released on May 29, 2019 (referred to as *T2T6 assembly*).

Figure 3 presents information about cenX assemblies. Since ONT assemblies often have inflated lengths of homonucleotide runs, we compressed each homonucleotide run into a single nucleotide (in the read set and assemblies) and recomputed the number of unique 19-mers. Figure 3 shows the distribution of frequencies of unique 19-mers in the compressed assemblies and illustrates that centroFlye and T2T6 assemblies have similar distributions of frequencies, while T2T4 assembly has significantly higher number of low-frequency unique 19-mers which are likely erroneous.

**Figure 3.**
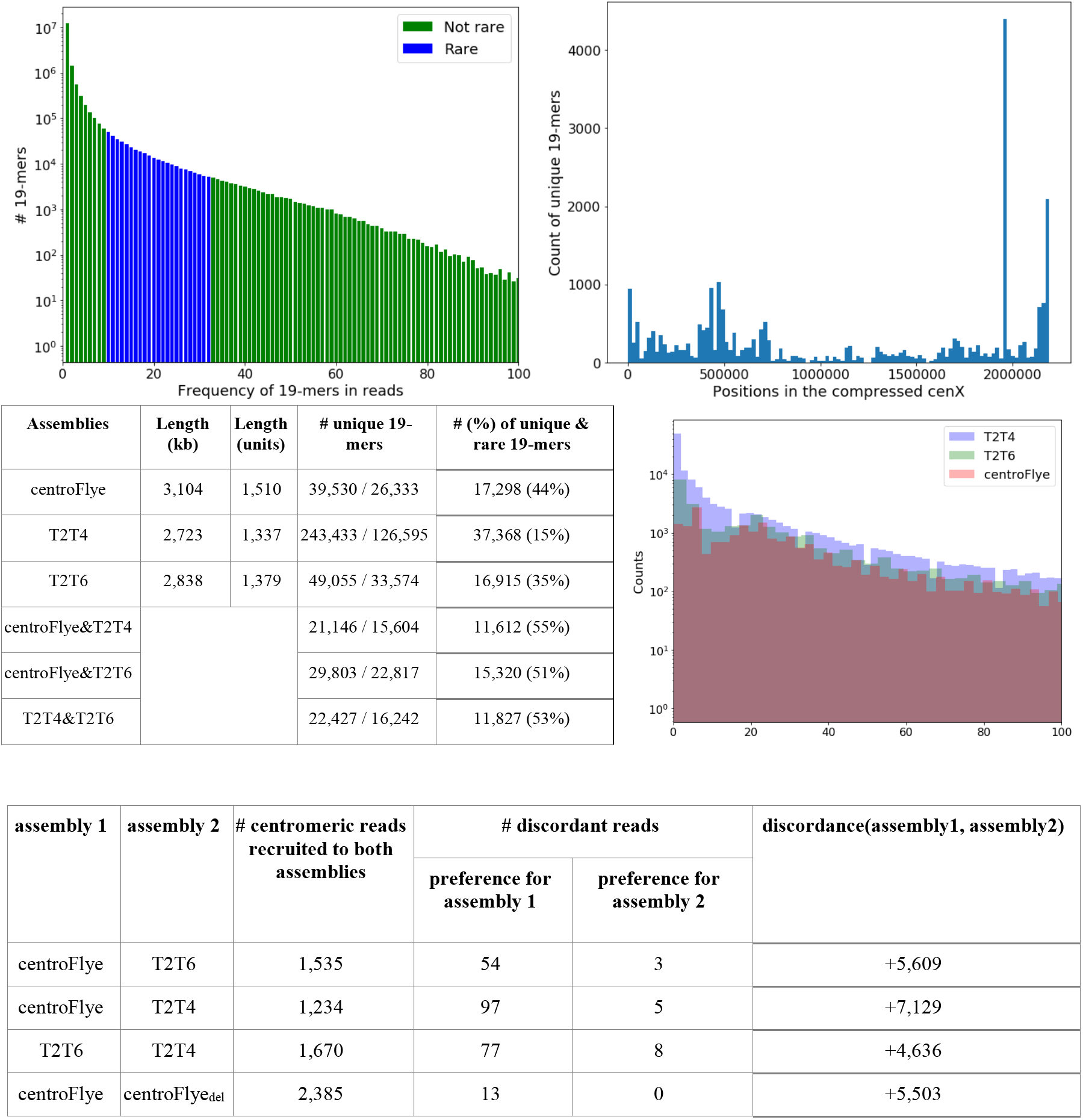
Information about cenX assemblies. **(Top left) Frequency histogram of 19-mers in the centromeric reads.** Each bar represents the number of 19-mers in units of centromeric reads with given frequency (log-scale). The bars corresponding to the rare 19-mers are shown in blue. Only 19-mers with frequencies that do not exceed 100 are shown. **(Top right) Distribution of unique 19-mers along the compressed cenX sequence.** Each bar represents the number of unique 19-mers in a segment of length ~20 kb (out of 26,724 unique 19-mers in the compressed cenX sequence). A large peak at positions 1,955,990 — 1,960,812 corresponds to a 4,822 nucleotide long (compressed) LINE insertion in cenX. **(Middle left) Comparison of centroFlye, T2T4, and T2T6 assemblies.** The column “number of unique 19-mers” shows the number of unique 19-mers before/after compression of homonucleotide runs. The column “number (percentage) of unique & rare 19-mers” refers to the number (percentage) of unique 19-mers in an assembly that are rare in reads. The T2T4 centromere has 57,811,689 — 60,534,892 coordinates and the T2T6 centromere has 57,827,622 — 60,665,308 coordinates on chromosome X. **(Middle right) Distribution of frequencies (in logarithmic scale) of unique 19-mers in the compressed centroFlye, T2T4, and T2T6 cenX assemblies. (Bottom) Number of recruited reads, discordant reads and the discordance score for all pairs of assemblies**.

### Mapping reads to assemblies

Given a centromere assembly, one can map all centromeric reads to this assembly using its unique *k*-mers. Below we describe how to use this mapping for comparison of various assemblies. To illustrate the effect of an indel on various quality metrics we introduced an artificial deletion of length 50 kb (25 units) in the centroFlye assembly at position 600 kb corresponding to 300 units (we refer to this artificial assembly as *centroFlye*_*del*_).

For each pair of assemblies, we used their shared unique 19-mers to align centromeric reads to them (an alignment is accepted if it includes at least three units and 20 unique 19-mers). Figure 4 compares positions of read alignments to each pair of assemblies and reveals structural discrepancies between them. Comparison of centroFlye and centroFlye_del_ assemblies reveals an expected discrepancy around unit 300. Both T2T4 and T2T6 assemblies differ from the centroFlye assembly around units 150-180 in the centroFlye assembly although the centroFlye and T2T6 are more complaint in this area. The size of deletion in T2T4 is approximately 32 units (around unit 135), and in T2T6 — only 5 units (around unit 175). Four other discrepancies are shared between both versions of T2T and the centroFlye assembly. They are located around the following units in the centroFlye assembly: 450 — 2 unit deletion in T2T4 (around unit 410) and 1 unit insertion in T2T6 (around unit 445), 750 — 56 units deletion in both T2T4 (around unit 700) and T2T6 assemblies (around unit 720), 1050 — 47 units deletion in both T2T4 (around unit 950) and T2T6 (around unit 975) assemblies, and 1400 — 36 and 24 units deletion in T2T4 (around unit 1,250) and T2T6 (around unit 1,300) respectively. Below we argue that T2T assemblies have putative misassembles in the surrounding areas.

**Figure 4.**
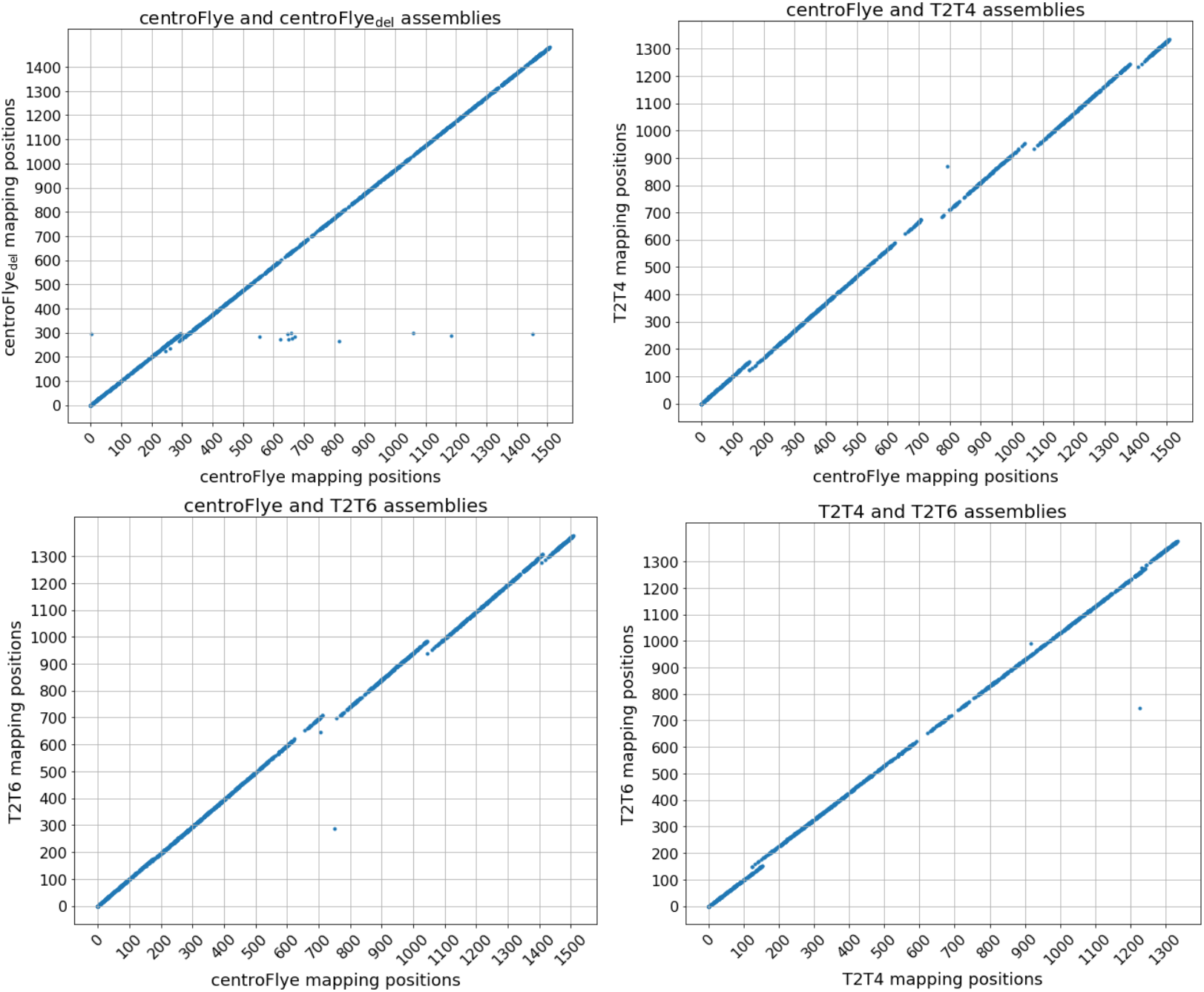
Comparison of read mappings between the centroFlye, centroFlye_del_, T2T4, and T2T6 assemblies. Each dot corresponds to a centromeric read. X- and Y-coordinate represent the starting unit position in the corresponding assemblies. (**Top Left**) Comparison of centroFlye and centroFlye_del_ assemblies reveals a discrepancy around unit 300. **(Top Right)** Comparison of centroFlye and T2T4 assemblies reveals discrepancies around units 150 (135), 450 (410), 750 (700), 1050 (950), and 1400 (1250) in the centroFlye (T2T4) assembly. **(Bottom Left)** Comparison of centroFlye and T2T6 assemblies reveals discrepancies around units 180 (175), 450 (445), 750 (720), 1050 (975), 1400 (1300) in the centroFlye (T2T6) assembly. **(Bottom Right)** Comparison of T2T4 and T2T6 assemblies reveals discrepancies around the unit 150 and 1240 in both assemblies.

### Quality assessment of the centromere assemblies

Although Appendix “Benchmarking centroFlye on simulated datasets” demonstrates that centroFlye accurately reconstructs simulated centromeres, it is unclear how to evaluate centroFlye assemblies of real centromeres. Benchmarking of various genome assemblers would not be possible without the quality assessment tools such as QUAST (Gurevich et al., 2013). However, since QUAST was not designed for analyzing long tandem repeats, it is not applicable for analyzing centromere assemblies. Below we describe some metrics for the reference-free quality assessment of centromere assemblies.

### Coverage test

Errors in an assembly affect the coverage near the assembly breakpoints. Thus, in the case of a uniform coverage, regions with abnormal coverage may point to assembly errors. For example, a deletion inflates the coverage near the deletion breakpoint (doubles the coverage in the case of a long deletion) and an insertion generally reduces the coverage in the inserted segment. Figure 5 shows the coverage plots for centroFlye, centroFlye_del_, T2T4, and T2T6 assemblies and reveals that a deletion in centroFlye_del_ assembly inflates the coverage by approximately 60% at the deletion breakpoint (from ~50x to ~80x).

**Figure 5.**
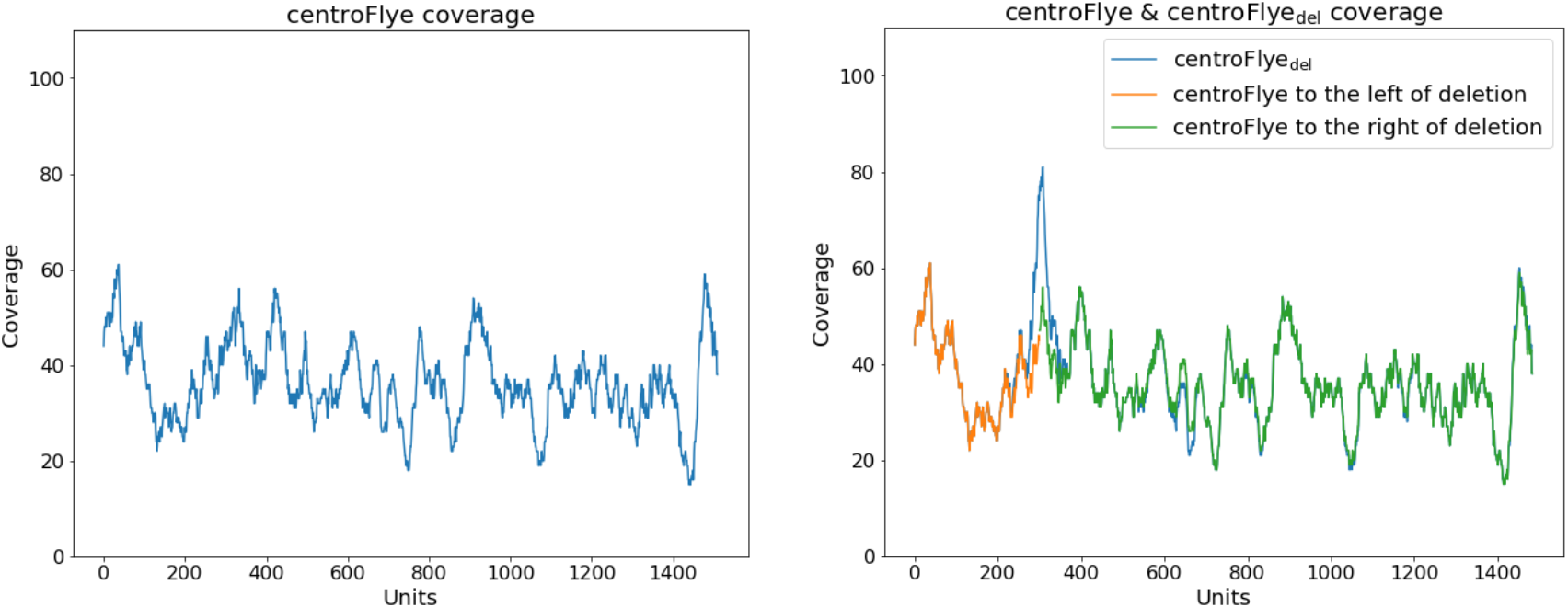
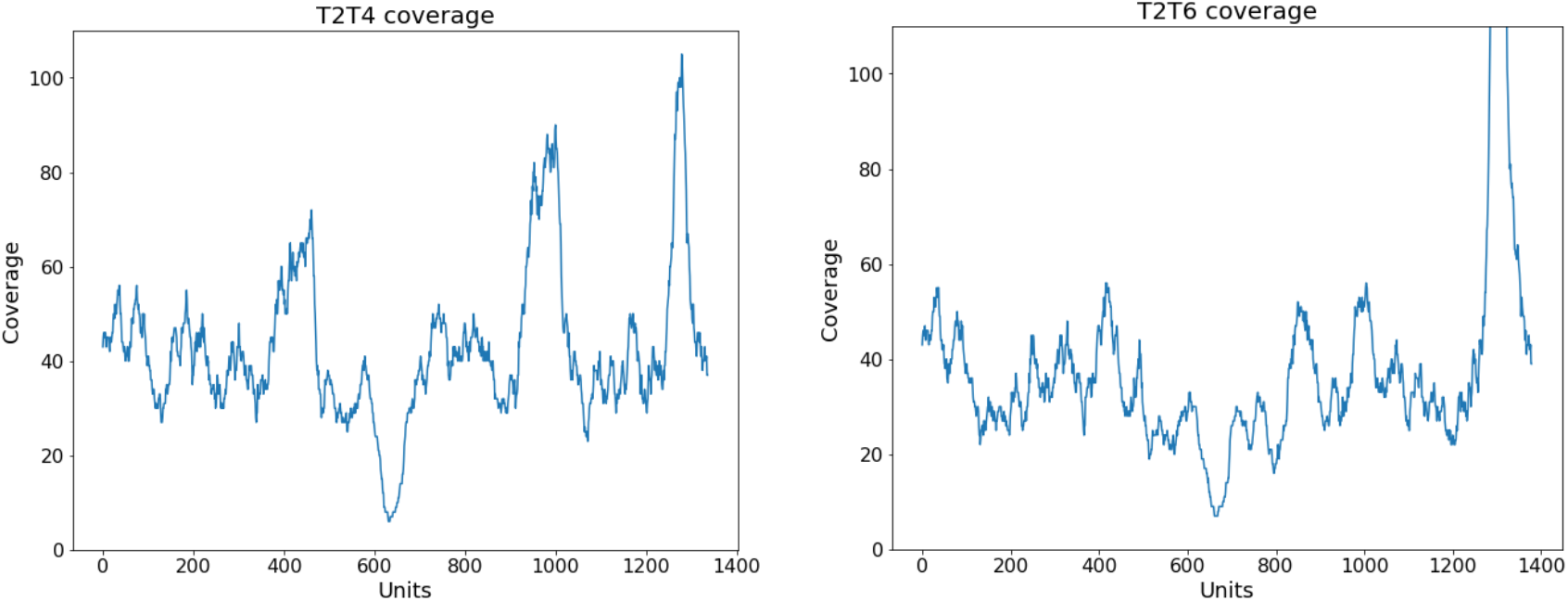
Coverage plots for centroFlye, centroFlye_del_, T2T4, and T2T6 assemblies. centroFlye_del_ refers to the centroFlye assembly of cenX with artificially introduced a 50 kb (25 units) deletion at position 600 kb (unit 300). The peak around unit 1,300 in T2T6 gives coverage almost 1,000 and was cut. Note, that the reduction in the number of mappable reads caused by the deletion in centroFlye_del_ assembly affects the coverage near the deletion in this assembly: 1,696 and 1,676 reads were mapped to centroFlye and centroFlye_del_ assemblies respectively.

Typical ONT datasets are characterized by a rather uniform read coverage, thus suggesting that such spikes in coverage may reveal assembly errors. However, it turned out that analyzed ONT dataset is characterized by a non-uniform read distribution, making it difficult to infer assembly errors from irregularities in the read coverage (see Appendix: “Analyzing non-uniform read coverage of the T2T4 read-set”). Even though T2T assemblies of cenX demonstrate higher coverage variations than the centroFlye assembly, these variations do not necessarily point to assembly errors, necessitating a need to introduce additional quality metrics for centromere assemblies. The T2T6 assembly coverage is more similar to the centroFlye assembly coverage (as compared to T2T4 assembly), but has a large spike at the end of the DXZ1 array, which is coordinated with a discrepancy in Figure 4.

### Discordance test

A *k*-mer is *shared* between an assembly and a read aligned to this assembly if it occurs in both the assembly and the read at the same position in their alignment. Given a set of *k*-mers *Anchors*, we define *shared*_*Anchors*_(*Read, Assembly*) as the number of *k*-mers from *Anchors* that are shared between *Read* and *Assembly*. The larger is *shared*_*Anchors*_(*Read, Assembly*), the better the assembly “explains” the read with respect to a given set of *k*-mers. Given a read-set *Reads*, we define *shared*_*Anchors*_(*Reads, Assembly*) as the sum of *shared*_*Anchors*_(*Read, Assembly*) over all reads in *Reads*.

To compare two assemblies, we define *Anchors* as the set of shared unique *k*-mers between them (the default value *k*=19) and compute the *discordance* between these assemblies as *discordance(Assembly’, Assembly’’)* = *shared*_*Anchors*_(*Reads, Assembly’*) – *shared*_*Anchors*_(*Reads, Assembly’’*). For centroFlye and T2T6 assemblies, *discordance*(centroFlye,T2T6) = 4,735, suggesting that the centroFlye assembly is a better fit for the read-set than the T2T6 assembly (Figure 3).

We classify a read *Read* as *discordant* with respect to assemblies *Assembly’* and *Assembly”* and a set of *k*-mers *Anchors* if there is a large difference (by at least *k*) between *shared*_*Anchors*_(*Read, Assembly’*) and *shared*_*Anchors*_(*Read, Assembly”*), thus showing preference for one of the assemblies. We say that a discordant read *votes* for *Assembly’* (*Assembly”*) if this difference is positive (negative). There are 54 (3) discordant reads voting for centroFlye (T2T6) assemblies. Figure 3 illustrates that the centroFlye assembly improves on all other assemblies with respect to the discordance scores.

A concentration of discordant reads at a certain region voting for *Assembly’* over *Assembly”* suggests that *Assembly”* has a multi-unit deletion in this region. Figure 6 reveals three clusters of discordant reads voting for centroFlye over T2T6 assembly at the regions ~200, ~400-600, ~1400 units in the centroFlye assembly (only two discordant reads vote for the T2T6 over the centroFlye assembly). These regions are coordinated with discrepancies shown in Figure 4 and likely point to large deletions in the T2T6 assembly. Similar comparison between centroFlye and T2T4 assemblies reveals putative misassembles in T2T4 at units ~200, ~400, ~800, ~1150, and ~1400 in the centroFlye assembly. As expected, comparison of centroFlye and centroFlye_del_ detects a single deletion at unit 300.

**Figure 6.**
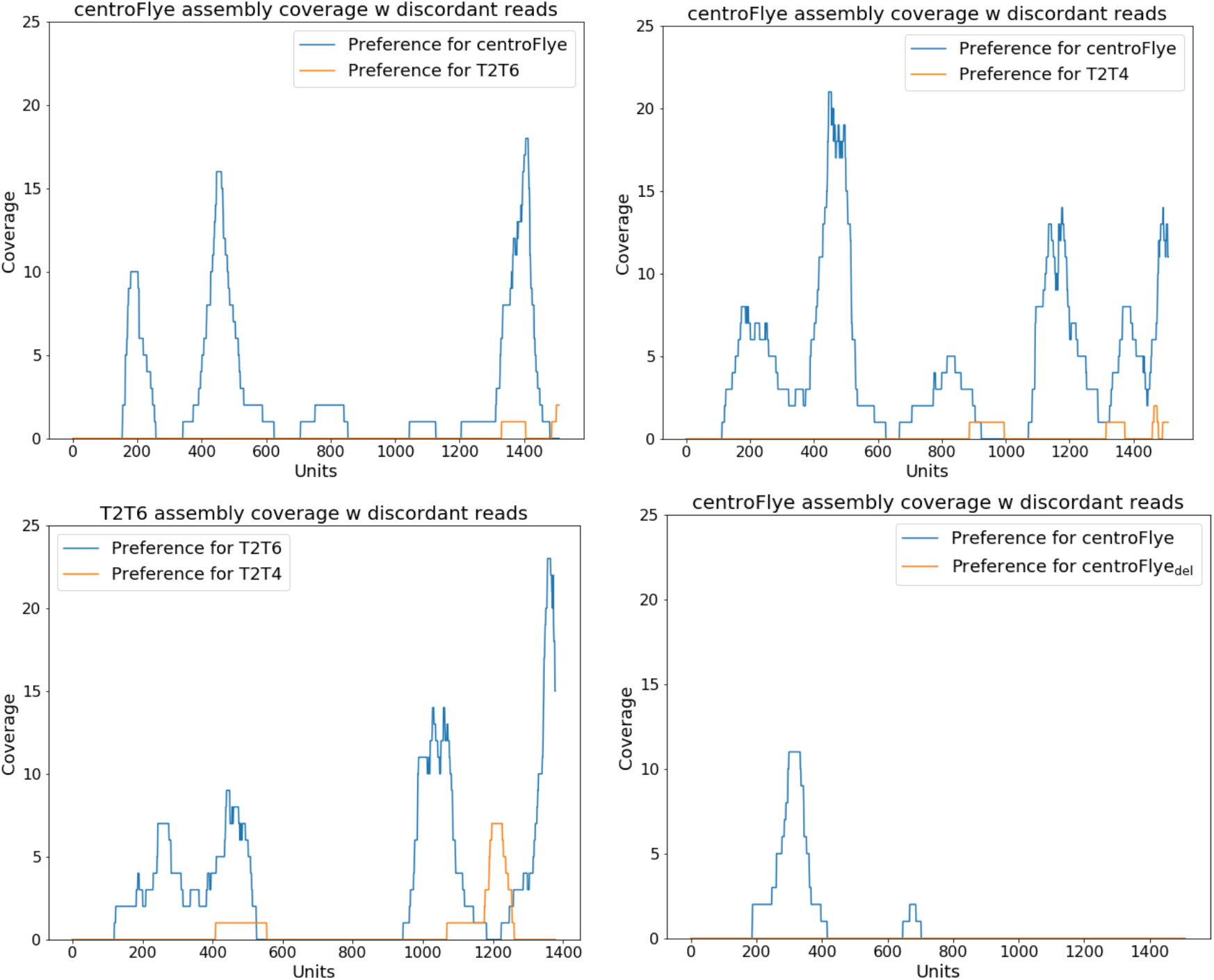
Coverage of various cenX assemblies by discordant reads.

Appendices “Hanging index test” and “Breakpoint test” introduce other assembly quality metrics and provide additional support for the centroFlye assembly.

### Variations in HORs provide insights into evolutionary history of cenX

Various copies of a repeat are usually partitioned into subfamilies that reflect evolutionary history of this repeat. For example, Jurka and Smith, 1988 discovered that Alu repeats in the human genome are split in AluJ and AluS subfamilies. Various follow-up studies increased the number of Alu subfamilies to over 200 and provided insights into evolutionary history of Alu repeats (Price et al., 2004). Some “active” copies of Alu repeats continue to produce variation by jumping to new genomic locations, thus posing the mutagenic threat to the human genome (Benett et al., 2008).

Since partitioning centromeric HORs into subfamilies remains unknown, it is difficult to reveal mutagenic units in centromeres and study their influence on disease. Appendix “Variations in HORs provide insights into evolutionary history of cenX” reveals subfamilies of the cenX HOR and demonstrates that their positions in cenX provide initial insights into the cenX evolution.

## Discussion

Although the role of centromeres (chromosome segregation) is conserved throughout evolution, the centromere sequences greatly differ across various species. For example, although X chromosome is highly conserved across all mammals, the mammalian X centromeres vary greatly across mammalian species. Since all previous attempts to develop an automatic tool for centromere sequencing have failed so far, centromeres remain the last frontier of genome sequencing and an enigma for both evolutionary and functional studies.

We compared centroFlye assembly to the manual T2T assemblies and identified several likely misassembles in the T2T6 assembly. Figure 4 reveals five large discrepancies (deletions) between centroFlye and T2T6 assemblies, Figure 5 shows highly inflated coverage around unit 1300 in T2T6 assembly, Figure 6 describes discordant reads that reveal three problems in T2T6 assembly, Figure J1 demonstrates high hanging index in five regions in T2T6 assembly, and Figure K1 suggests five potential breakpoints in the T2T6 assembly.

Although centroFlye opens a possibility to fill the largest remaining gaps in the human genome and study centromere evolution, further algorithmic developments are needed to sequence all human centromeres. We are currently working on the following tasks to accomplish this goal:

### Extending centroFlye for reconstructing multi-HOR centromeres

Although some centromeres are characterized by a single HOR (like DXZ1 HOR on chromosome X or D16Z2 HOR on chromosome 16), others contain multiple HORs. For example, chromosome 3 is built from four HORs: D3Z1 (total length 2102 kb on cen3), D3-2 (total length 461 kb), and two recently identified HORs with total lengths 217 kb and 64 kb, respectively (Uralsky et al., 2019). To modify centroFlye for assembling multi-HOR centromeres, we need to extend the classification of reads. For example, instead of the current classification into prefix, internal, and suffix centromeric reads, we need to introduce D3Z1/D3-2 reads (that start from D3Z1 repeats and end with D3-2 repeats) among many other read-types.

### Extending centroFlye for reconstructing centromeres with shared HORs

Some chromosomes share the same HOR with other chromosomes, for example human chromosomes 1, 5, and 19 share the same HOR referred to as D1Z7/D5Z2/D19Z3 (Uralsky et al., 2019). centroFlye faces additional complications in such cases since the centromeric reads that align to D1Z7/D5Z2/D19Z3 will sample three chromosomes rather than a single one. In this case, instead of classification into prefix, internal, and suffix centromeric reads, we need to classify reads into three types of prefix reads (for each of chromosomes 1, 5, and 9), internal reads, and three types of suffix reads.

### Extending centroFlye for analyzing abnormal HORs

Although centroFlye revealed 37 abnormal HORs in cenX assembly, these HORs were not used to guide the assembly (Appendix: “Abnormal units in centroFlye cenX assembly”). We plan to detect abnormal HORs in reads and to incorporate them in the centroFlye algorithm.

### Extending centroFlye into tandemFlye for analyzing arbitrary tandem repeats and resolving them in the assembly graph

centroFlye currently requires a HOR and a read-set as an input. However, after three decades of HOR sequencing, new HORs are still being discovered (Uralsky et al., 2019) and it remains unclear how accurate the currently known HORs are. Our goal is to discover new HORs in a *de novo* fashion and to develop a tandemFlye tool for reconstructing arbitrary tandem repeats. We plan to further incorporate tandemFlye into the tandem repeat resolution unit in Flye by automatically deriving a set of HORs for each chromosome.

### Extending centroFlye for reconstructing centromeres in diploid genomes

Although centroFlye reconstructed the human haploid X centromere, reconstructing diploid centromeres represents a difficult algorithmic challenge.

## Methods

### Recruiting centromeric reads (Figure 2.1)

centroFlye recruits centromeric reads for a specific chromosome by identifying all reads that align to HORs from this chromosome. It uses the *fitting alignment* (Šošić and Šikić, 2017) of HORs to all reads and recruits reads with sequence identity exceeding a threshold. centroFlye uses a *sequence identity threshold* (the default value is 83% because most human HORs differ from the HOR consensus by less than 5% and the error rate in reads is ≈12%). In the case of the human X centromere, centroFlye recruits 2,680 centromeric reads (total length ~133 Mb) that align to DXZ1 or its reverse complement.

Below we assume that all centromeric reads have DXZ1 in the forward orientation and complement a read if it is not the case (see Appendix: “Analysis of reads with reported DXZ1 alignments to both strand orientations”). 150 centromeric reads have lengths varying from 2 to 5 kb, 1,382 reads are longer than 30kb and 897 of them are ultralong. The longest centromeric read is 527 kb.

### Partitioning centromeric reads into units (Figure 2.2)

DXZ1 was derived at the dawn of the sequencing era based on limited sequencing data (Waye and Willard, 1985). Appendix: “Deriving accurate consensus HORs” describes how to infer a more accurate consensus HOR for a centromere. This approach revealed a new DXZ1* consensus of length 2054 that differs from DXZ1 by 30 substitutions and indels (~1.5% difference).

We use the Noise-Cancelling Repeat Finder (NCRF) (Harris et al., 2019) to partition each centromeric read into units. Given a read and a consensus HOR, NCRF partitions a read into units, each unit representing a single copy of a HOR. Although NCRF was not designed to characterize the possible gaps between units (e.g., transposon insertions or small rearrangements), this limitation of NCRF does not significantly affect the centroFlye results. If NCRF reports several alignments for a given read, the longest one is kept. Incomplete units appearing at prefix or suffix of a read are discarded.

centroFlye discards all centromeric reads with (the longest) alignment shorter than three units. 295 centromeric reads (including 42 ultralong reads) of total length 7 Mb were discarded. NCRF identified 56,138 units in the remaining 2,385 centromeric reads, including 39,363 units in the remaining 855 ultralong reads.

### Classifying centromeric reads into prefix, internal, and suffix reads (Figure 2.3)

A centromeric read is classified as a prefix (suffix) read if it has a prefix (suffix) of length at least *prefixThreshold* that does not match the HOR consensus (default threshold *prefixThreshold*=50 kb). NCRF revealed 15 prefix, 2,357 internal, and 13 suffix reads. This classification is important for “moving inside the centromere” using an approach similar to the approach for reconstructing unbridged repeats in Flye (Kolmogorov et al., 2019).

### Identifying rare centromeric *k*-mers (Figure 2.4)

We define the *frequency* of a *k*-mer as the number of occurrences of this *k*-mer in units of reads from a centromeric read-set. centroFlye identifies rare centromeric *k*-mers by analyzing *k*-mers with frequencies that fall into a predefined interval (centroFlye uses the default value *k*=19). Since a HOR may be repeated in a centromere thousands of times (with small variations), we expect most *k*-mer from a HOR to have high frequencies (Figure I1). Our goal is to identify *rare k*-mers that appear just once (*unique k*-mers) or a few times in a centromere and use them for centromere assembly. Note that a *k*-mer from a genome “survives” without errors in a read with the *survival rate* that can be approximated as *survivalRate*(*k*)=(1-*p*)^*k*^, where *p* is the probability of an error at a given position of a read. A more accurate estimate from the real data suggests that the survival rate of 19-mers in the ONT reads is 0.34. Thus, since the recruited set of ultralong cenX reads has ≈32x coverage, we expect that a unique *k*-mer from a given position in the genome survives in ≈11 ultralong centromeric reads.

We define the interval [*bottom, top*] and classify a *k*-mer as rare if its frequency is larger than *bottom*survivalRate*coverage* and smaller than *top*survivalRate*coverage* (the default values *bottom*=1 and *top*=3). Although 391,361 *k*-mers from centromeric reads were classified as rare (Figure 3, top left), many of them represent erroneous versions of *k*-mers from a HOR copy rather than truly rare *k*-mers in cenX. Indeed, the number of reads containing a given *k*-mer *b* is affected by the number of genomic positions with *k*-mers similar to *b* since error-prone reads covering these similar *k*-mers may contain *b*. Since many HOR copies may contain *k*-mers similar to a unique *k*-mer, this observation explains the complications in inferring the set of unique/rare *k*-mers. For example, a single nucleotide insertion in a *k*-mer from a genome occurs in an ONT read with probability 0.03 (Lin et al., 2016). Thus, each such insertion in a *k*-mer from DXZ1* has a high chance to be classified as a rare *k*-mer. Below we describe how to filter out such spurious rare *k*-mers using the distance graph.

### Constructing the distance graph (Figure 2.5)

The key observation to separate false positives from unique *k*-mers is that the distances between unique *k*-mers in reads are likely to be conserved but distances between false positive *k*-mers are not necessarily conserved. The distance graph reveals pairs of unique *k*-mers that are separated by roughly the same distance in multiple reads.

Given a set of rare centromeric *k*-mers *V*, we define the weighted directed *distance graph* with the vertex-set *V* and the edge set defined as follows. Two vertices *v* and *w* are connected by a directed edge of length *ℓ* > 0 if there is a centromeric read where *w* follows *v* at the distance *ℓ* units. If there are *t* different edges between *v* and *w* of the same length *ℓ*, we combine them into a single *multiedge* (*v, w, ℓ*) of multiplicity *t*. We further remove all multiedges with multiplicities below *minCoverage = C*survivalRate*coverage* (the default value *C*=0.4 and, thus, *minCoverage*=4 for our set of rare centromeric reads). Finally, we remove all *conflicting* parallel multiedges from the graph (multiedges connecting the same vertices but having different lengths) and all isolated vertices. The remaining vertices form a set of only 28,703 *k*-mers out of 391,361 initially constructed rare centromeric *k*-mers (below we refer to the remaining *k*-mers as “unique”). Even though, some of the remaining *k*-mers turned out to be rare rather than unique (see Appendix: “Analysis of frequencies of recruited *k*-mers in the centroFlye cenX assembly”), they still appear to be valuable for assembly efforts.

### Reconstructing the centromere (Figure 2.6)

centroFlye reconstructs the centromere sequence using an approach similar to the approach for resolving unbridged repeats in Flye (Kolmogorov et al., 2019). Instead of using the *divergent positions* (like in Flye), it uses the unique centromeric *k*-mers to iteratively reconstruct the centromere.

Given a unit in a read, a *cloud* of this unit is defined as the set of unique *k*-mers occurring in this unit. centroFlye represents each read as a sequence of clouds (*cloud-sequence*) that we refer to as *readCloud*. Given two clouds *c* and *c’*, we define *shared(c,c’)* as the number of shared unique *k*-mers in these clouds. Given two cloud-sequences of the same length *x*=*x*_1_, … *x*_*n*_ and *y*=*y*_1_, … *y*_*n*_, their score is computed as Σ_*i=1,n*_ *shared*(*x*_*i*_,*y*_*i*_). Given cloud-sequences *x* of length *n* and *y* of length *m*, their *i-score* (1≤ *i*≤ *n*) is defined as the score between *x*_*i*_, … *x*_*min(i+m,n)*_ and the prefix *y*_1_, …*y*_*min(m,n-i+1)*_ of *y.* The *maxScore* between *x* and *y* is defined as the maximum of *i*-scores over all possible values of *i* and an alignment of *x* and *y* that achieves the *maxScore* value is referred to as an *optimal alignment*.

An alignment of multiple reads (a *contig*) defines an alignment of their cloud-sequences — a cloud-sequence that we refer to as *cloud-contig* (for multiple aligned units in this alignment, their combined cloud is defined as the union of all individual clouds). centroFlye supports an operation of optimally aligning a new read against a cloud-contig and updating the cloud-contig to include a newly added read.

Figure 2.6 illustrates the centroFlye repeat resolution algorithm. First, all prefix reads are aligned based on their prefixes, represented as cloud-sequences and combined into an initial cloud-contig that starts at the unit position 0. Afterwards, centroFlye selects a still unaligned read with a highest-scoring optimal alignment against the cloud-contig (in case of ties, the read with the rightmost starting position of the optimal alignment is selected). If the score of this alignment exceeds *stopThreshold*, centroFlye adds this read to the growing contig-cloud, otherwise it stops the contig extension (the default value *stopThreshold* = 10). In case centroFlye incorporates nearly all reads in the growing contig-sequence (including suffix reads), we classify the centromere construction as successful and proceed to the polishing step (see Appendix: “Analysis of centromeric reads that do not map to centroFlye assembly”). Otherwise, we apply the same centromere construction procedure but this time starting from suffix rather than prefix reads. It may happen that the prefix-based centromere construction stops before completing the centromere but the suffix-based construction generates the entire centromere. If both prefix-based and suffix-based centromere reconstructions stop, we generate the suffix and prefix contig-sequences that do not span the entire centromere.

Note that at this step we do not obtain the cenX sequence, but rather the cloud-sequence of the centromere and the starting unit positions inside the cenX for all aligned centromere reads.

### Polishing the reconstructed centromere sequence (Figure 2.7)

Using reported starting unit positions for all centromeric reads, centroFlye separately polishes each HOR unit of the (yet unknown) centromere separately. In the case of cenX, it selects unit of the median length from corresponding reads, uses it as a template, and applies four rounds of the polishing algorithm in Flye (Lin et al., 2016, version 2.5).

Polishing results in a sequence of length 3,103,541 that includes 1510 units and a single insertion of a LINE repeat (see Appendix: “Search for Alu insertions in cenX”). It contains 39,530 unique 19-mers, i.e., 19-mers that appear only once in the assembly. Since ONT assemblies have high rates of homonucleotide indels, we further compressed all homonucleotide runs in the polished centromere, resulting in a *compressed centromere* that has only 26,333 unique 19-mers (Figure 3, top right).

## Acknowledgements

We are indebted to Ivan Alexandrov, Mikhail Kolmogorov, Karen Miga, and Valery Shepelev for many insightful comments that improved centroFlye algorithm. We are also grateful to Anton Bankevich, Alexander Bzikadze, Alla Mikheenko, Adam Phillippy, Cynthia Wu, and Jeffrey Yuan for helpful discussions and suggestions.

## Author Contributions

All authors contributed to developing centroFlye algorithm and writing the paper. A.B. implemented centroFlye algorithm. P.A.P. directed the work.

## Code availability

Main codebase of the algorithm is available at https://github.com/seryrzu/centroFlye. Jupyter notebooks for reproducing the figures in this study are provided in the Github repository https://github.com/seryrzu/centroFlye_paper_scripts. centroFlye centromere X assembly and all supporting data is available at Zenodo: https://doi.org/10.5281/zenodo.3369553.

## Appendix A. Analysis of centromeric reads that do not map to centroFlye assembly.

centroFlye assembly contains 39,530 unique 19-mers (including 17,298 rare 19-mers) that we used to align 2,385 centromeric reads that contain at least 3 units according to NCRF. 689 (~29%) reads were unmapped with total length 15.3 Mb (~4.9x coverage of cenX). 13 unmapped reads were longer than 100kb. However, NCRF did not report a long continuous alignment for these reads, suggesting that they are either recruited in error or have a very high error-rate. Total length of 1,696 mapped reads is 110,739,155 and, thus, they correspond to ~35.7x coverage of cenX. Figure A1 shows distributions of lengths of mapped and unmapped reads.

For a read *Read*, we define *K-merPerUnitRatio(Read)* as the average number of unique 19-mer in the centroFlye assembly per unit in *Read*. The difference between *K-merPerUnitRatio* for unmapped and mapped reads in the centroFlye assembly is significant (two-sided Mann-Whitney U test, p-value < 10^−5^) with mean of *K-merPerUnitRatio* 46 and 57 for unmapped and mapped reads, respectively. However, if we consider only 17,298 rare 19-mers in these reads, *K-merPerUnitRatio* are 2.3 and 7.8 for unmapped and mapped reads respectively (same test, p-value < 10^−5^).

**Figure A1.**
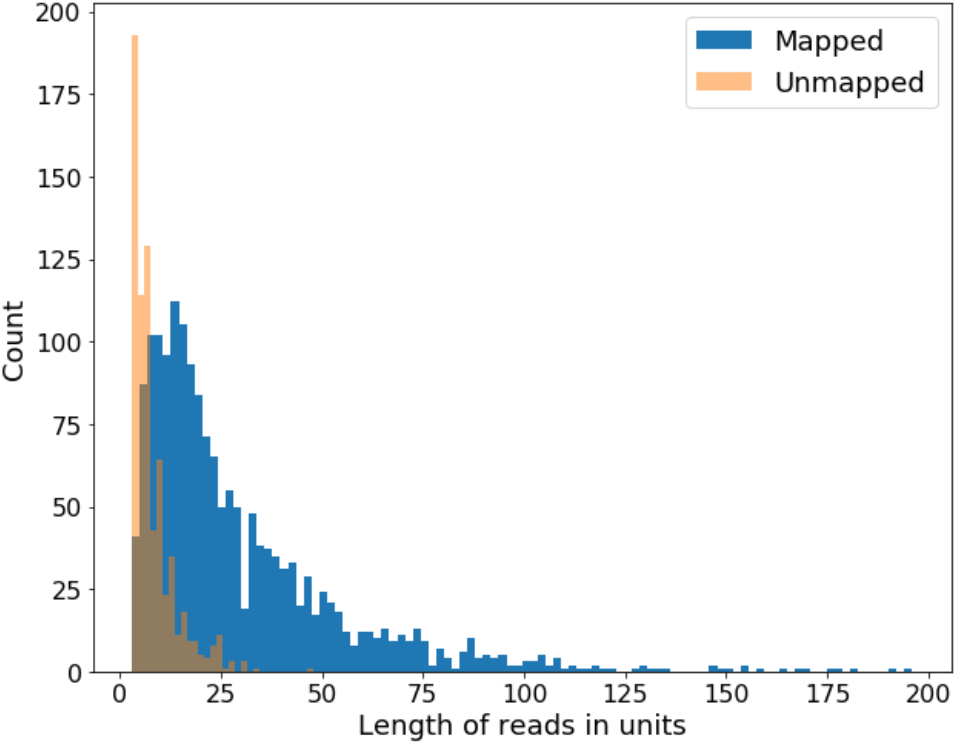
Distribution of lengths of mapped and unmapped reads. Read lengths are calculated in units.

## Appendix B. Analysis of reads with reported DXZ1 alignments to both strand orientations

We discovered only 4 centromeric reads (out of 2,385) that have alignments to DXZ1 in both strand orientations of length at least one unit (Figure B1). Considering average coverage of the dataset we conclude that these reads are likely to be chimeric, implying that there are no inversions of length longer than one unit on cenX.

**Figure B1.**
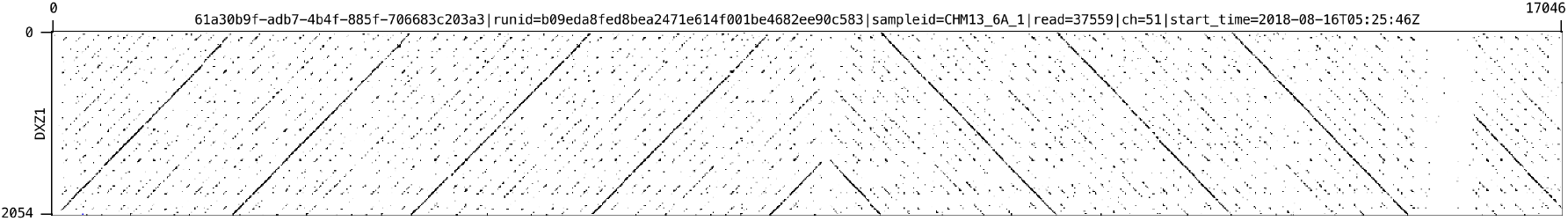
Dot-plot of a read with alignments to DXZ1 in both strand orientations. X-axis is the read, Y-axis is DXZ1.

## Appendix C. Search for Alu insertions in cenX

For all 2,680 centromeric reads (including 2,385 with DXZ1 alignments of length at least 3 units) we examined regions spanning from the start of leftmost DXZ1 alignment to the end of rightmost DXZ1 alignment as reported by NCRF (minimal alignment length is 500 nucleotides). We used the fitting alignment (Šošić and Šikić, 2017) of all members in the Alu repeat family (RepBase database, version 24.10) to the selected regions of reads with a predefined threshold (70%). We did not identify any selected regions in reads that contain Alu repeats. However, we identified 27 reads that contain Alu repeat units outside of the selected regions. 1 read was classified as a prefix read, 5 others — as suffix reads. 14 reads were unmapped. This result suggests that DXZ1 array in cenX does not contain Alu repeats.

## Appendix D. Benchmarking centroFlye on simulated datasets

In order to benchmark centroFlye, we simulated a tandem repeat of length ~1.03Mb which is a concatenation of 500 randomly mutated copies of DXZ1* that diverge from the consensus of DXZ1* by *Divergence* % (substitutions only). centroFlye successfully reconstructed tandem repeats for *Divergence* = 1%, 0.5%, and 0.1%. However, it failed to incorporate all internal reads to the cloud contig (Figure 2.6) for extremely low *Divergence* = 0.01%. Below we describe results on simulated tandem repeat with *Divergence* = 1%. Total number of substitutions in the simulated tandem repeat — 10,390, number of unique 19-mers — 52,738. We have added randomly generated flanking sequences of length 200kb on each side of the tandem repeat.

### Simulated reads

NanoSim (Yang et al., 2017) was trained on a subset of the first 1 million reads from the real T2T dataset with <u>T2T assembly v0.4</u> used as the reference with the following command:

~~~
~/soft/NanoSim/src/read_analysis.py -i rel2_1mm.fasta -r
../../../assemblies/chm13.draft_v0.4.fasta -t 32
~~~

The trained model was used to simulate 3000 reads (-n 3000) from the flanked repeat. 1150 reads were simulated as “unaligned” (see https://github.com/bcgsc/NanoSim) and are useless for assembly. The total length of 1850 “aligned” reads (without their low-quality ends) is 60,796,193 and, thus, the average coverage of the tandem repeat is ~43x. NCRF reported alignments to DXZ1* for 1380 reads. 1147 of those reads had (longest) alignment longer than 3 units.

### Unique 19-mers

140,780 19-mers were classified as rare and 34,717 were recruited as “unique”, 31,927 (92%) of which were unique in the tandem repeat. Only 2,790 (8%) recruited 19-mers were not unique in the tandem repeat. 1,547 of these k-mers are absent from the assembly, 1,242 appear 2 times in the assembly, and 1 appears 3 times.

### Reconstructed sequence

We applied centroFlye to the simulated reads in the same way it was applied to cenX assembly. 13, 1127, and 7 reads were classified as prefix, internal, and suffix reads, respectively. Only 1 out of 1147 reads was not mapped to the assembly. The length of reconstructed assembly is 500 units and coincides with the length of the simulated tandem repeat. Error-rate of the polished assembly is 0.7% after 4 iterations of polishing (edit distance 7281 between reconstructed and real tandem repeat sequences).

## Appendix E. Analyzing non-uniform read coverage of the T2T read-set

Median coverage of the whole chrX in HG38 with ultra-long reads (longer than 50 kb) is 30x, 5-percentile is 19x and 95-percentile is 43x. We applied moving average to the coverage with window 10,000. The median length of regions with coverage higher than 43x or lower than 19x is 23,893 (mean is 10,237). Figure E1 shows a surprisingly non-uniform coverage over chrX in HG38 with ultra-long reads (longer than 50 kb).

**Figure E1.**
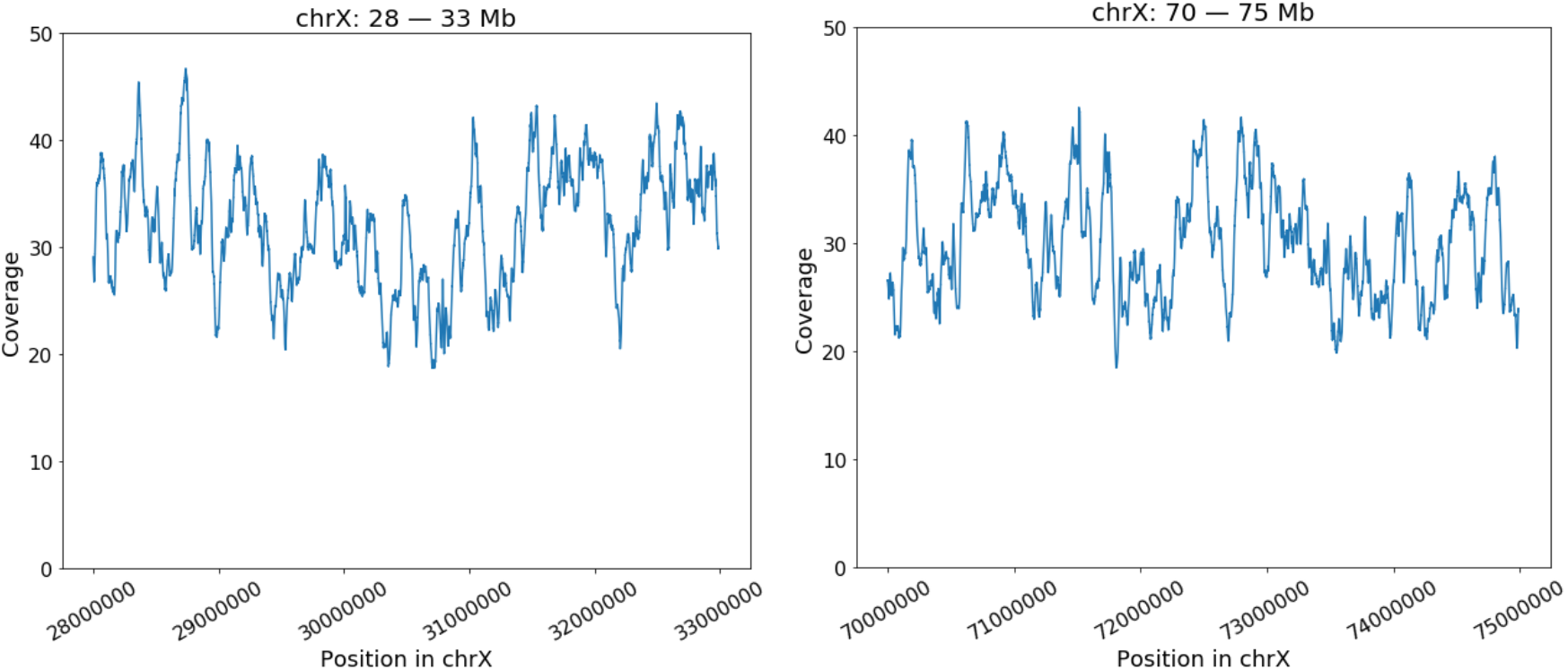
Coverage of chrX in HG38 with ultra-long reads. Range of coverage is [0, 50]. Moving average with window 10,000 in the region 28 — 33 Mb (**Left)** and the region 70 — 75Mb (**Right**).

## Appendix F. Abnormal units in centroFlye cenX assembly

Figure F1 presents the distribution of unit lengths in the centroFlye assembly of cenX and reveals that the vast majority of them have lengths similar to the length of the canonical DXZ1 HOR (the median unit length is 2057 and the standard deviation is 84). However, the length of 35 units differs from the average unit length by at least 5%, suggesting that such units may represent abnormal HORs rather than the canonical 12-mer DXZ1.

Table F1 presents dot-plots of 35 units with abnormally low/high unit lengths. Units with positions 0 and 1509 are the flanking units in the assembly. Majority of other units (except unit 334, cell 2×1 in the Table F1) have a HOR structure that is a concatenation of a prefix and a suffix of DXZ1*. Such units can be both longer or shorter than the canonical 12-mer DXZ1*. For example, a 17-mer unit at position 548 (cell 3×1) consists of a 9-mer prefix and 8-mer suffix while a 10-mer unit at position 624 (cell 4×2) consists of a 4-mer prefix and 6-mer suffix. Prefix or suffix can be absent (units 1347 and 1350). Unit 334 has a more complex structure and is a concatenation of monomers 8 — 10, 5 — 12, and 1 — 5.

We further aligned each unit against DXZ1* and analyzed units with large edit distance (greater than 100) in order to identify abnormal 12-mer units (of regular length) that however have a different sequence of monomers than DXZ1*. This analysis revealed two units — 333 and 335 (Figure F2). Unit 333 presents a circular shift of DXZ1 starting from the eighth alpha-monomer. Unit 335 is a concatenation of monomers 6-9 and 5-12. Table F1 and Figure F2 suggest that the region between units 332 and 335 contains multiple structural variations. Figure F3 summarizes the partitioning of this region into units provided by NCRF.

Table F2 summarizes the distribution of unique *k*-mers from the centroFlye assembly in each of the abnormal units and illustrates that 13 abnormal units don’t have any unique 19-mers in them. This observation suggests that these abnormal units are under-utilized in the cenX assembly. In the future, we plan to improve centromere assembly by incorporating both unique *k*-mers and abnormal units in the centroFlye algorithm.

**Figure F1.**
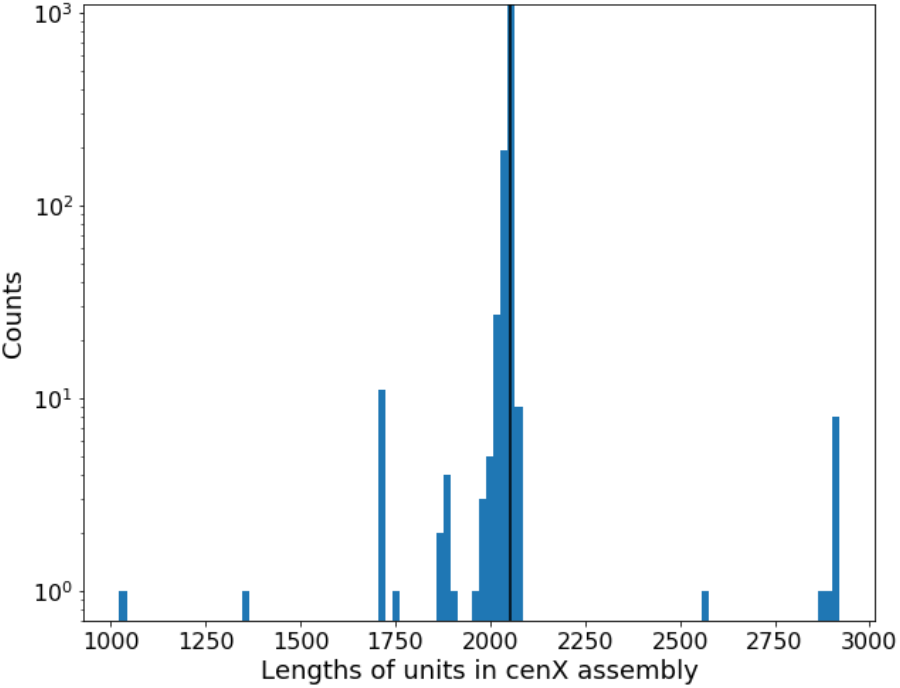
Distribution of unit lengths in the centroFlye assembly of cenX. Black vertical line shows the length of DXZ1* = 2054 nucleotides.

**Table F1:**
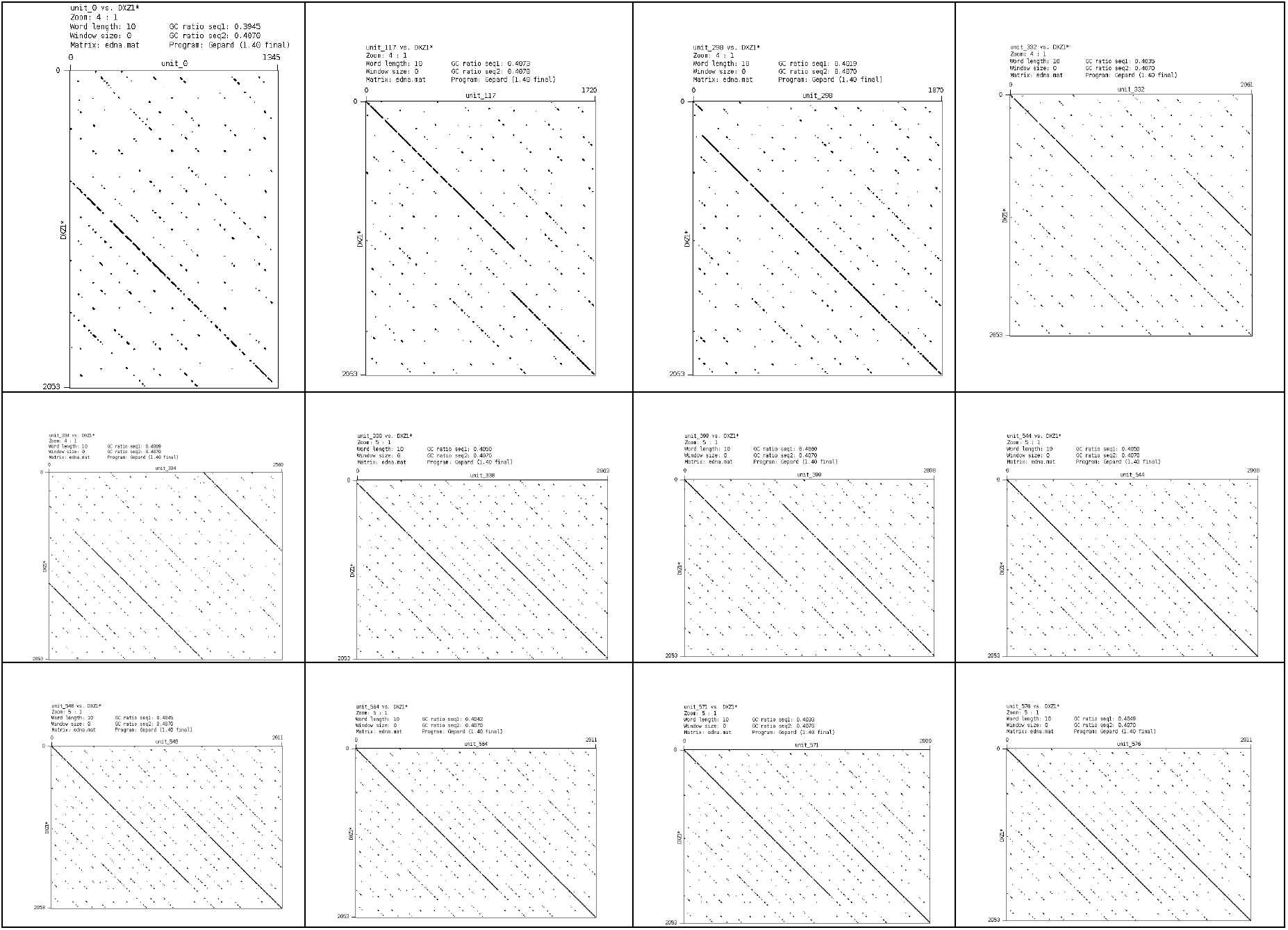
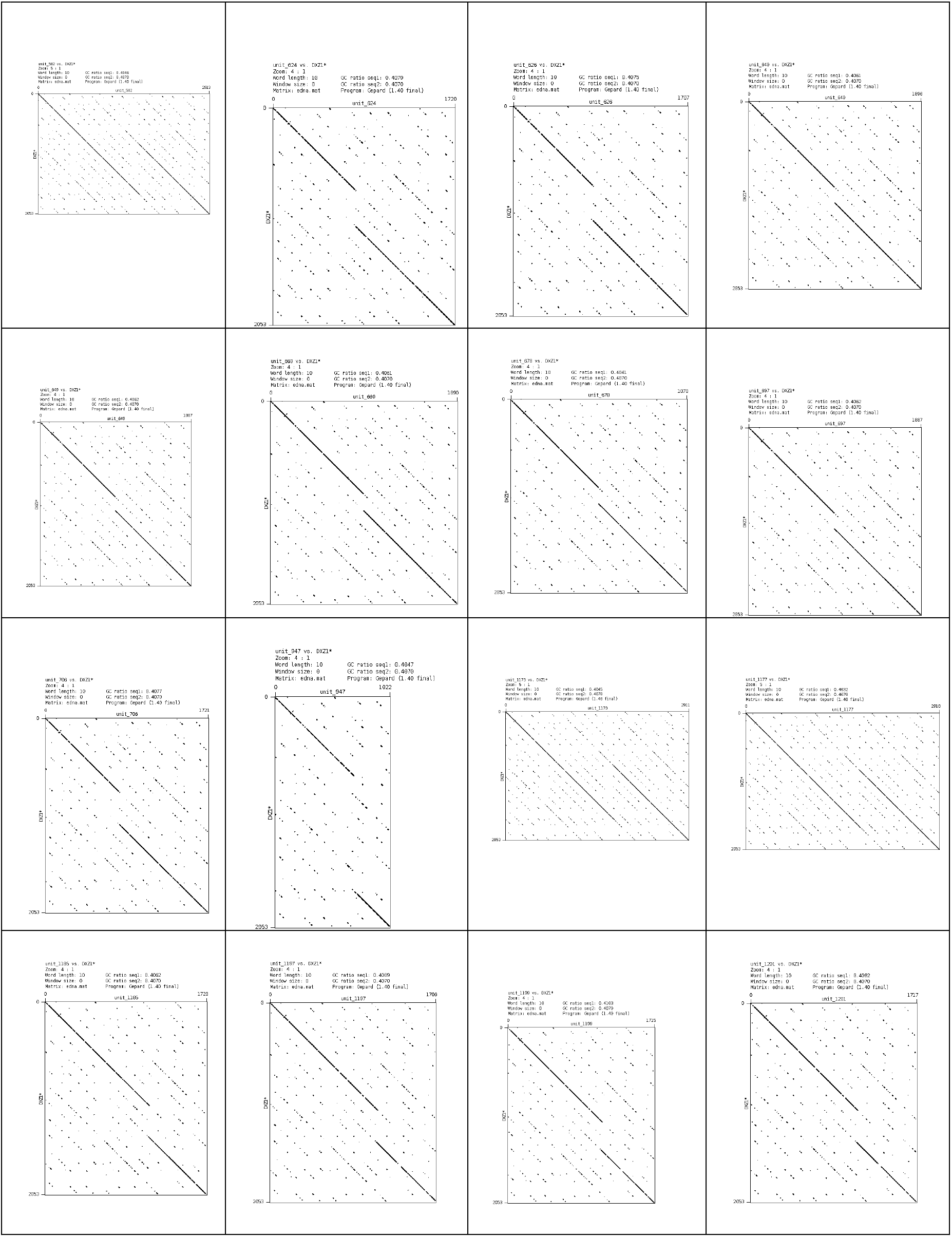
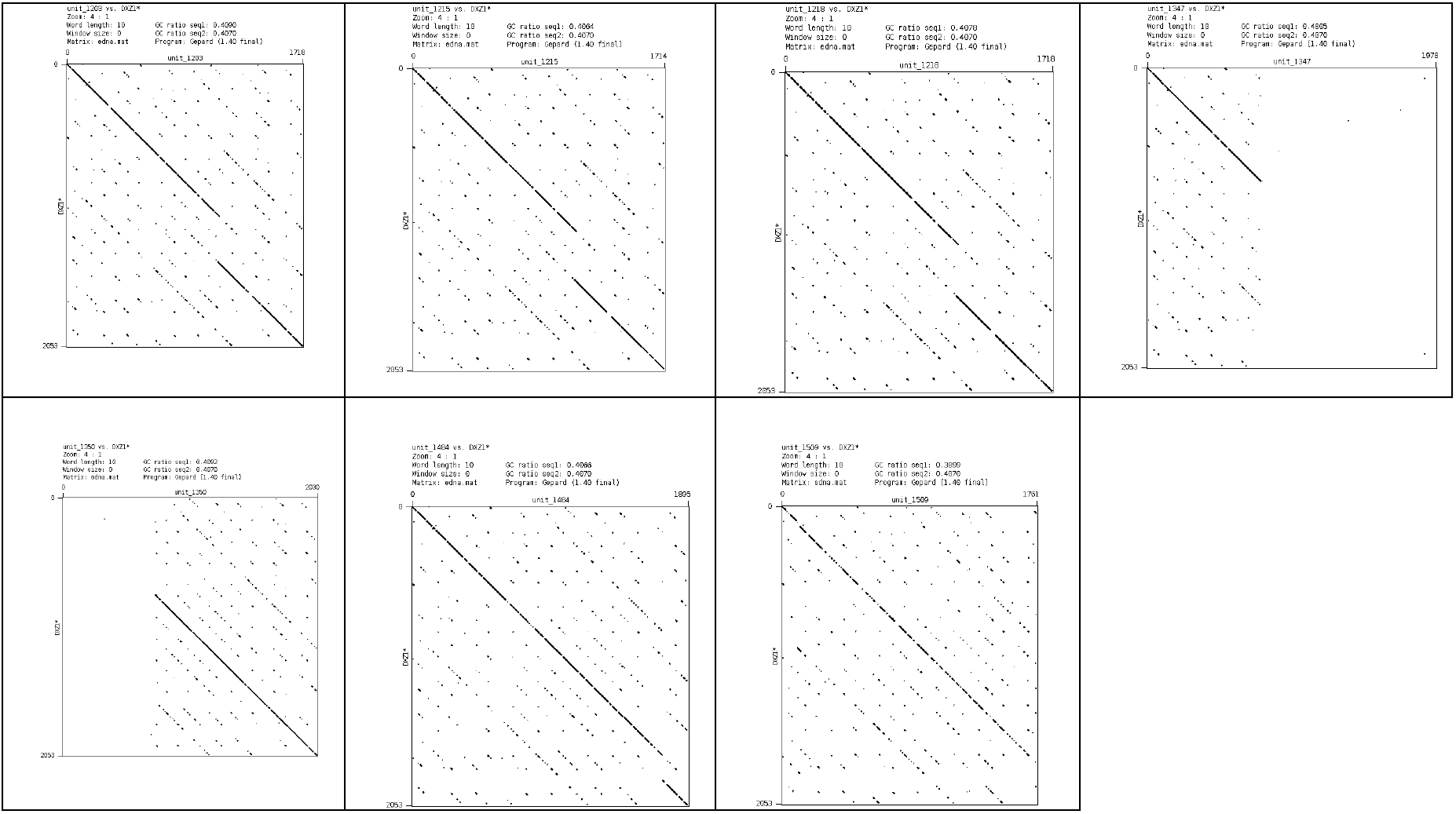
Dot-plots of 35 abnormal units against DXZ1*. X-axis corresponds to the unit. Y-axis corresponds to DXZ1*. The unit number is given in the X-axis label of each dot plot. Units are sorted in increasing order of their position in the cenX assembly and distributed by row. The shortest unit is a 5-mer (unit 1347) and a longest unit is a 17-mer (10 such units). The abnormal unit numbers (lengths) are 0 (8 units), 117 (10 units), 298 (11 units), 332 (11 units), 334 (15 units), 338 (17 units), 399 (17 units), 544 (17 units), 548 (17 units), 564 (17 units), 571 (17 units), 576 (17 units), 582 (17 units), 624 (10 units), 626 (10 units), 640 (11 units), 649 (11 units), 669 (11 units), 678 (11 units), 697 (11 units), 706 (10 units), 947 (6 units), 1173 (17 units), 1177 (17 units), 1185 (10 units), 1197 (10 units), 1199 (10 units), 1201 (10 units), 1203 (10 units), 1215 (10 units), 1218 (10 units), 1347 (5 units), 1350 (8 units), 1484 (11 units), and 1509 (10 units). Note a large cluster of closely located long 17-mer units (shown in blue) and two large clusters of closely located short 10-mer and 11-mer units (shown in green).

**Figure F2:**
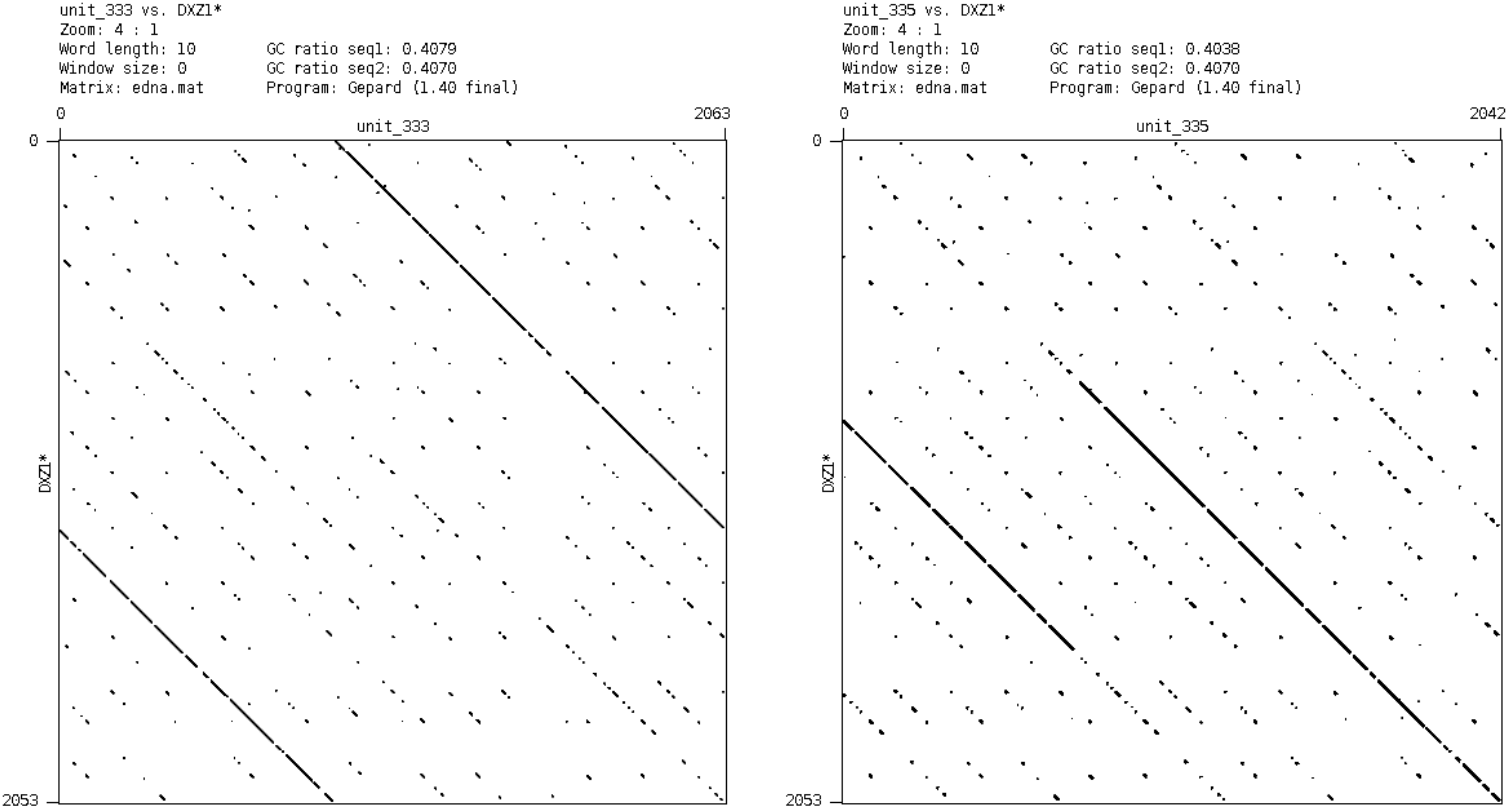
Dot-plots of units 333 (left) and 335 (right) against DXZ1*.

**Figure F3:**
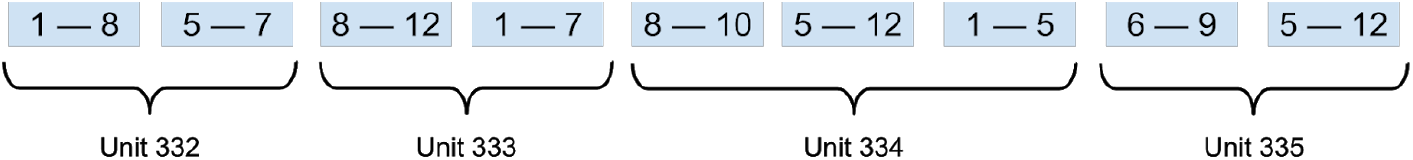
Partitioning of the region between units 332 and 335 into monomers.

**Table F2.**
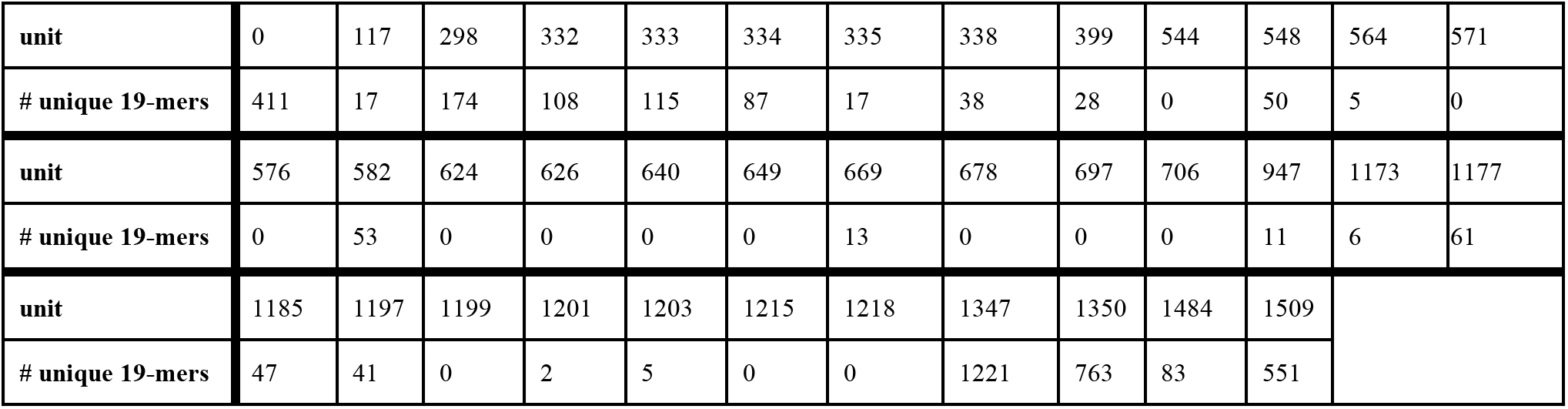
Number of unique 19-mers in each abnormal unit in the cenX assembly.

## Appendix G. Analysis of frequencies of recruited *k*-mers in the centroFlye cenX assembly

centroFlye cenX assembly contains 39,827 unique 19-mers. centroFlye recruited 28,703 of putative unique 19-mers from ONT reads. 16,488 (57.4%) of these 19-mers are also unique in the polished assembly. 9,267 (32.3%) are absent from the assembly and only 2,948 (10.3%) are repetitive in the assembly. Figure G1 presents a bar plot with frequencies in the assembly of the recruited 19-mers.

**Figure G1:**
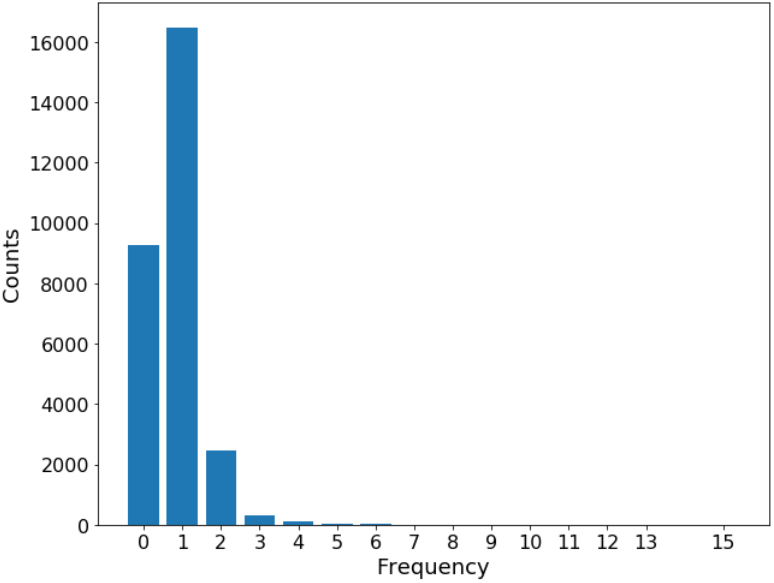
Frequencies of the recruited 19-mers in the centroFlye cenX assembly.

## Appendix H. Variations in HORs provide insights into evolutionary history of cenX

Below we reveal subfamilies of the cenX HOR and demonstrate that their positions in cenX provide initial insights into the cenX evolution.

To reveal subfamilies of the cenX HOR, we align each standard unit in homonucleotide-compressed centroFlye cenX assembly (1471 out of 1510 units) to the compressed DXZ1* and use the resulting multiple alignment to construct the *profile logo* for every position of DXZ1*. Figure H1 illustrates that the vast majority of positions in units are highly conserved (average percent identity 98.8% when considering deletions as mutations and 99.5% when excluding them, Figure H2), some positions reveal significant variations. We select *divergent* positions such that the ratio of the *secondary vote* to the *majority vote* is greater than *minRatio* (default value *minRatio* = 0.3) and the *secondary vote* is not a deletion. This procedure reveals 7 divergent positions. Each divergent position is characterized by its *majority vote* frequency and *secondary vote* frequency

We consider pairs of divergent positions that are at least *minDistance* (default value *minDistance* = 100) positions apart and select the “most correlated” pairs using the *biprofile* approach from Keich and Pevzner, 2002. Similarly to Price et al., 2004, we apply the *χ^2^* test of independence and select the pair of divergent positions with the smallest *p*-value. The divergent positions 580 and 695 with majority votes 71% and 72% (for nucleotides ‘AA’) and secondary votes 30% and 28% (for nucleotides ‘GG’) were selected. For these positions, the biprofile frequency for ‘AA’ and ‘GG’ is 69% and 27%, respectively, resulting in p-value < 10^−10^. We split the set of all units into two clusters with respect to nucleotides at the selected pair of positions, resulting in a split of all standard units into two clusters with 1010 and 389 units, respectively. We repeat this process iteratively for each of the new clusters until no more clusters of size greater than *minSize* (default value *minSize* = 100) are generated. This process stops after two iterations and results in four subfamilies of cenX HOR. Figure H3 shows the distribution of units in these subfamilies along the cenX assembly: red and green subfamilies are located in the middle and the second half of the assembly (~units 500-1450), orange subfamily is concentrated in the beginning (~units 1-300), blue subfamily — between green and orange (~units 300-550). Note that units in different subfamilies are alternating along cenX, providing an initial insight into how subfamilies “colonize” cenX during its evolution.

Since we derived HOR subfamilies without using positions of the units, clustering of each subfamily into close positions along cenX suggests that the algorithm originally described for identifying Alu subfamilies (Price et al., 2004) also works for HOR subfamilies. However, our results also reveal a newly observed phenomenon of *HOR recombination* and suggest that analysis of centromere evolution may be more complex that analysis of Alu evolution (Figure H4).

**Figure H1.**
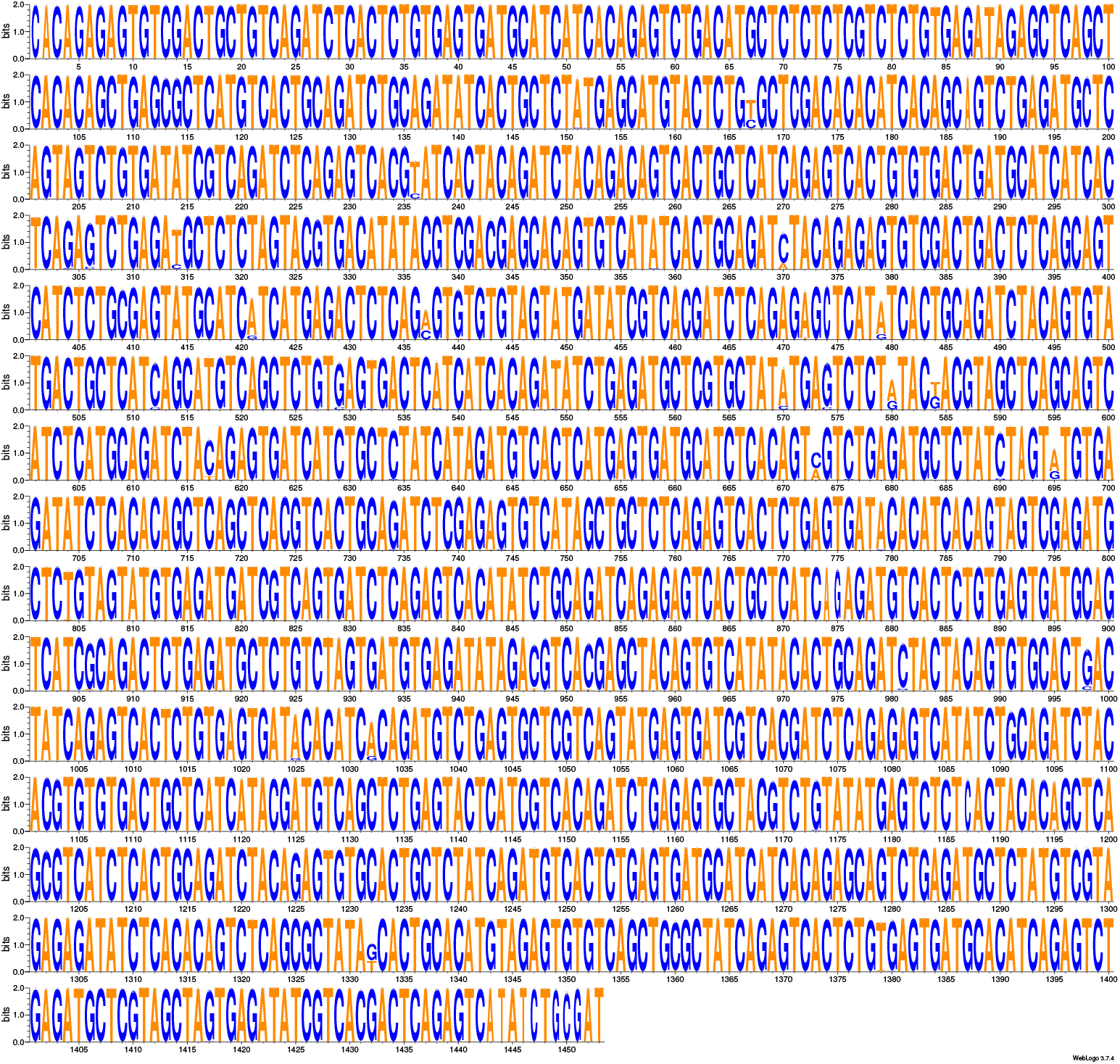
Profile logo of the alignment of 1472 compressed units to the compressed DXZ1*. Plot generated by WebLogo (Crooks et al, 2004). “Narrow” letters (for example, positions 874, 875) represent positions with high frequency of a deletion. Although most positions are highly conserved, there exist 7 divergent positions: 167, 437, 580, 584, 673, 695, and 1332.

**Figure H2:**
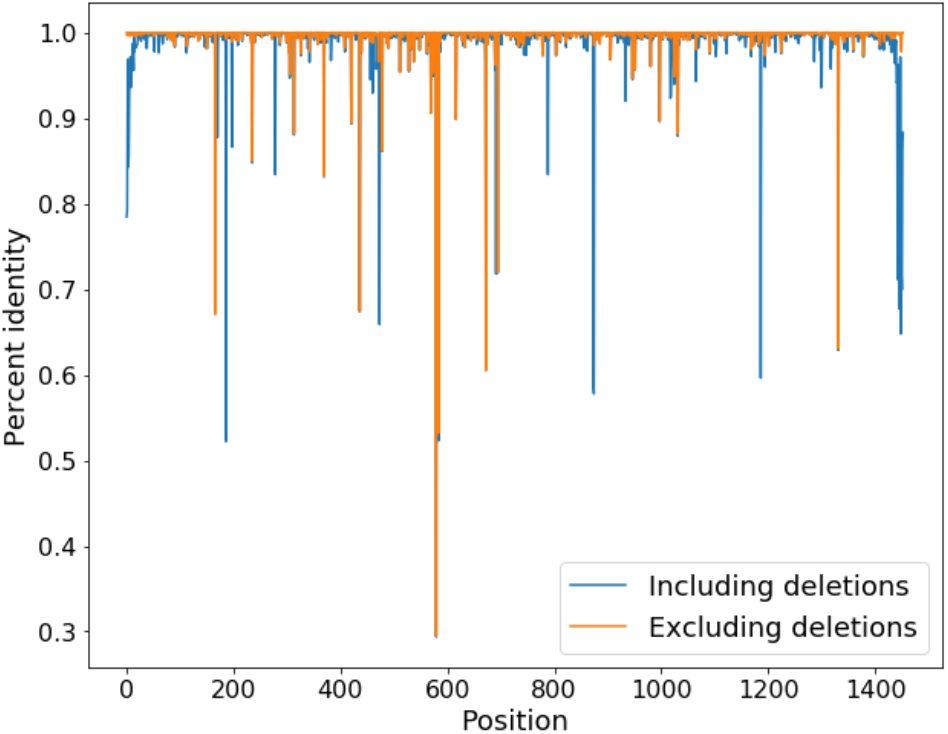
Percent identity for alignment of compressed DXZ1* to the compressed units in cenX assembly. When considering deletions as mutations: mean percent identity = 98.8%, median = 99.8%. When ignoring deletions: mean percent identity = 99.5%, median = 100%.

**Figure H3:**
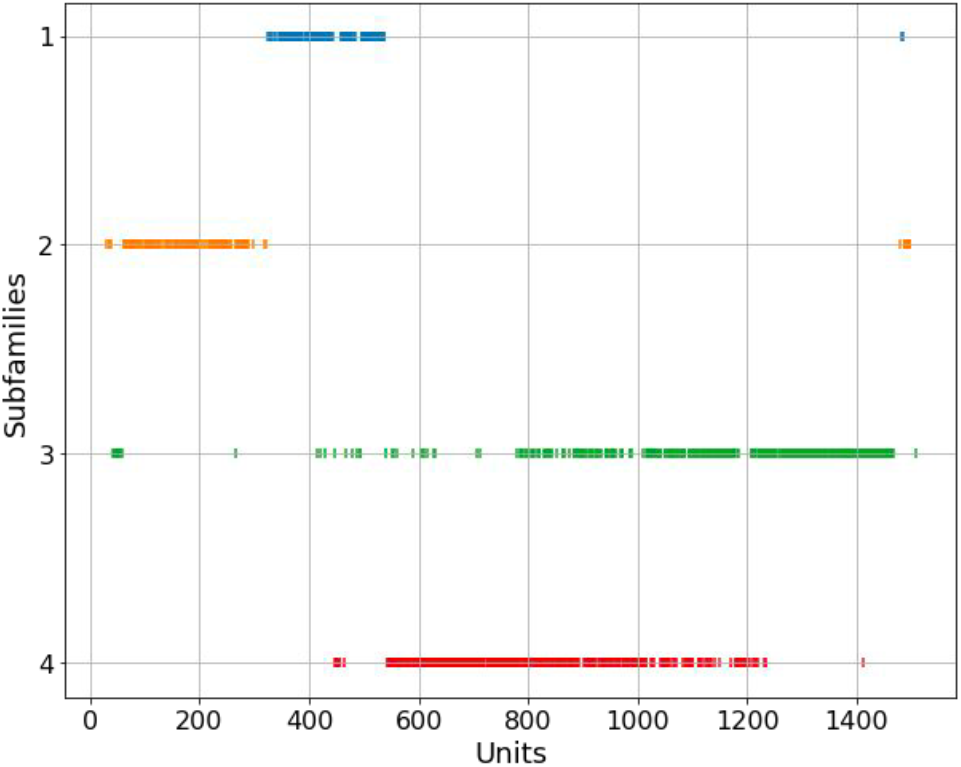
Distribution of four subfamilies of cenX HOR along cenX (*minSize* = 100). Red, green, orange, and blue subfamilies have 389, 429, 140, and 132 units.

**Figure H4.**
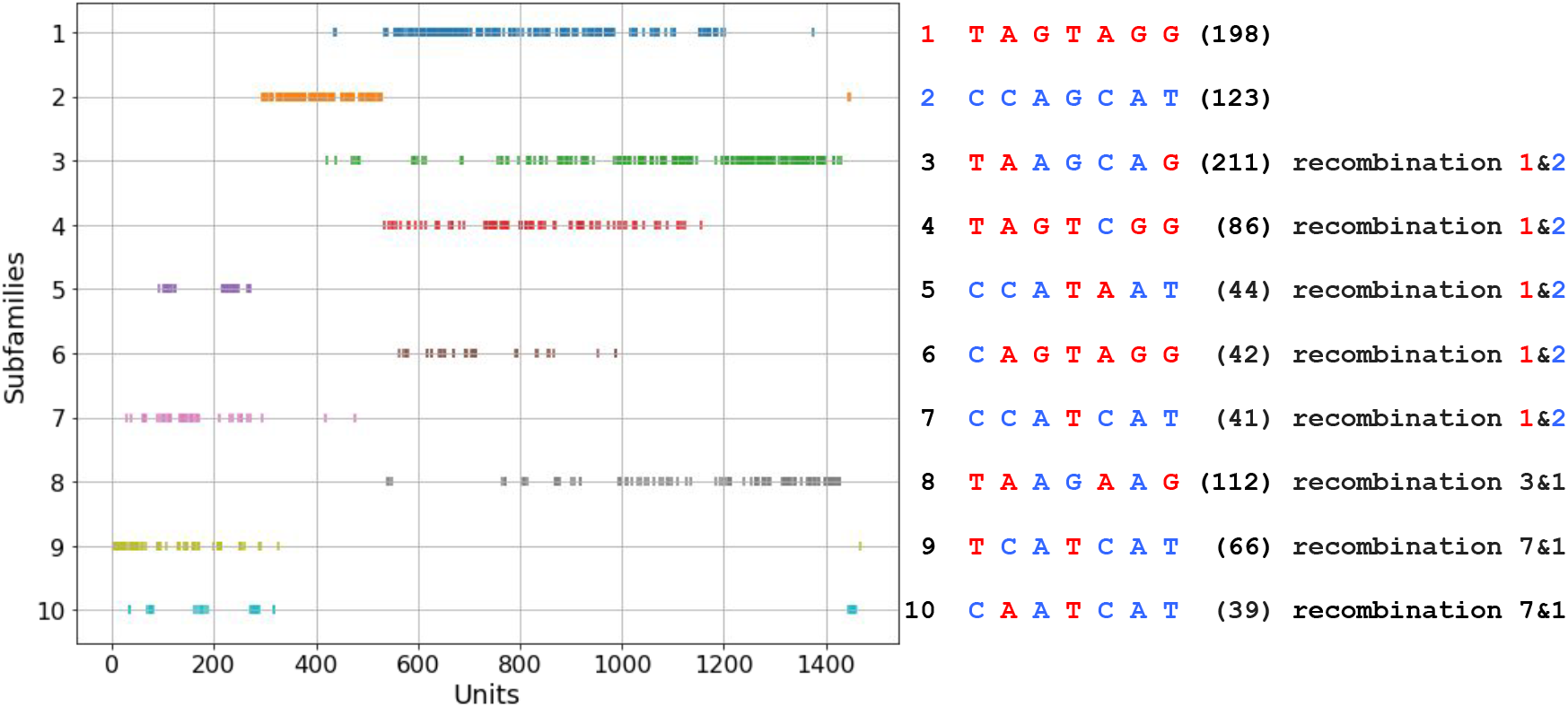
HOR recombination. With exception of only eight units, each of seven divergent positions has only two nucleotides in each of the standard units: T or C (position 167), A or C (position 437), G or A (position 580), T or G (position 584), A or C (position 673), G or A (position 695), and G or T (position 1332). We exclude these eight units from consideration and classify the remaining units into 2^7^= 64 clusters depending on the string that they spell out in seven divergent positions. Two of these strings (the “red” string TAGTAGG and the “blue” string CCAGCAT) differ at each position. Ten largest clusters (shown above along with the number of units in each cluster) correspond to strings that represent recombination events between the red and the blue strings (string 3-7) or between the red string and a recombinant of red and blue strings (strings 8-10). The ancestral human cenX HOR is likely located within positions 500-1200 occupied by clusters 1, 4 and 6 with HORs that are most similar to the gorilla cenX HOR (Durfy and Willard, 1990).

## Appendix I: Deriving accurate consensus HORs

Figure I1 reveals that some 19-mers from DXZ1 have surprisingly low frequencies in centromeric reads (234 of them have frequencies below 100), raising a concern that DXZ1 deviates from the consensus of all copies of the HOR on the X chromosome. To derive a more accurate consensus of a HOR, we define the *frequency* of a circular sequence as the minimum frequency of its *k*-mers (with respect to a given read-set). Given a set of centromeric reads *Reads*, we define its consensus HOR as a circular sequence (of length at least *L*) with maximum frequency with respect to this read-set among all circular sequences (if there are multiple such sequences, we select the shortest one). Thus, finding a consensus HOR is reduced to finding a long directed cycle in the de Bruijn graph *DB*_*k,t*_(*Reads*) of *k*-mers with frequencies exceeding *t* (we use the default parameters *L*=2000 and *k*=30 and found the maximum *t* such that *DB*_*k,t*_(*Reads*) has a long directed cycle). This approach revealed a new DXZ1* consensus of length 2054 that differs from DXZ1 by 30 substitutions and indels (~1.5% difference, Figure I1).

**Figure I1.**
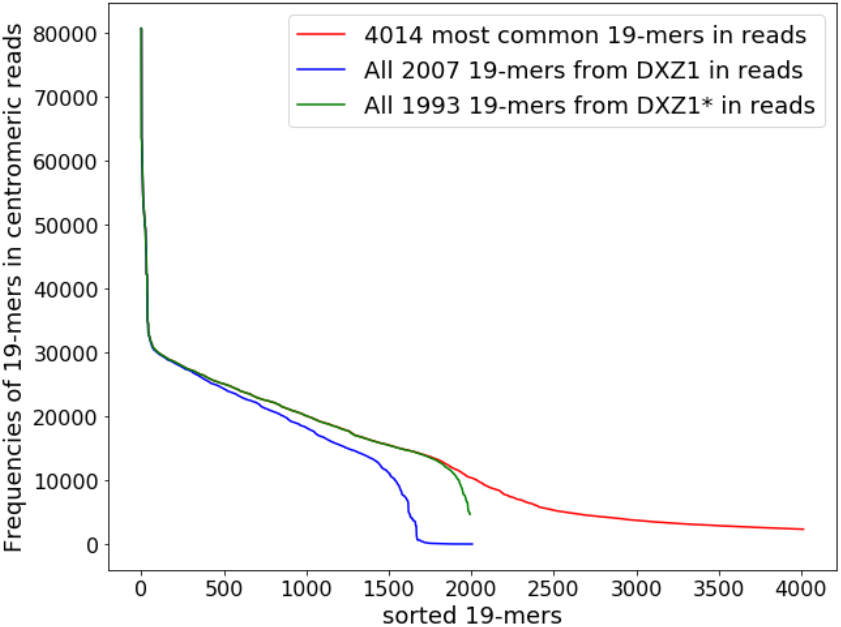
Frequent 19-mers in centromeric reads. For each curve, all 19-mers are ordered along the x-axis in the decreasing order of their frequencies. The blue and green curves represent frequencies of all 2,007 19-mers from the circularized DXZ1 and DXZ1*, respectively (DXZ1* is a more accurate consensus HOR for cenX). The red curve represents the frequencies of 2 * 2,007 = 4,014 most frequent 19-mers from all centromeric reads.

## Appendix J: Hanging index test

Consider an ultralong (longer than 50 kb) read *Read* spanning units from *a* to *b* in a centromere and a unit *a* < *x* < *b.* We compare the set of identified unique *k*-mers appearing in the segment [*a, a+l*] of the centromere with the set of unique *k*-mers appearing in the first *l* units of *Read* and say that *Read* has a *hanging prefix* of length *l* if these sets do not overlap. For each read, we define its *longest hanging prefix* (*longest hanging suffix* is defined similarly) and limit attention to reads that have hanging prefixes (suffixes) of length at least *buffer* units (the default value *buffer* = 2). For each unit in the centromere that has at least *minUniqueKmers* unique *k*-mers (default value *minUniqueKmers* = 5) we define its *hanging prefix (suffix) coverage* as the number of hanging prefixes (suffixes) in reads that span this unit. Finally, we compute the *hangingCoverage* of a unit as the average of its hanging prefix coverage and hanging suffix coverage and define the *hangingIndex* as the ratio of *hangingCoverage* and the total coverage of a unit. Units with high *hangingIndex* point to potential misassembles (the default threshold for *hangingIndex* is 0.15). We compute moving average of *hangingIndex* with the window 5 units.

The mean values of the *hangingIndex* for centroFlye, centroFlye_del_, T2T4, and T2T6 assemblies are 0.019, 0.024, 0.058, and 0.031, respectively (Figure J1). The hanging index test reveals an artificial deletion in centroFlye_del_ at unit 300 (*hangingIndex* 0.19). The hanging index test reveals several putative deletions in the T2T4 assembly (around units 150, 350 and 500-650, 1000, and 1250). We also detect several putative misassembles in the T2T6 assembly — around units 180, 450, 650-750, 1000 and 1300 with hanging indexes 0.4, 0.22, 0.18, 0.17 and 0.22 respectively. Locations of these potential misassembles are largely coordinated with Figure 4 and Figure 6. It is likely that the T2T6 assembly may have breakpoints at units 180, 450, 650-750, 1000, and 1300 potentially caused by multi-unit indels. These regions seem to provide a challenge for various assembly efforts and are likely to be misassembled.

**Figure J1.**
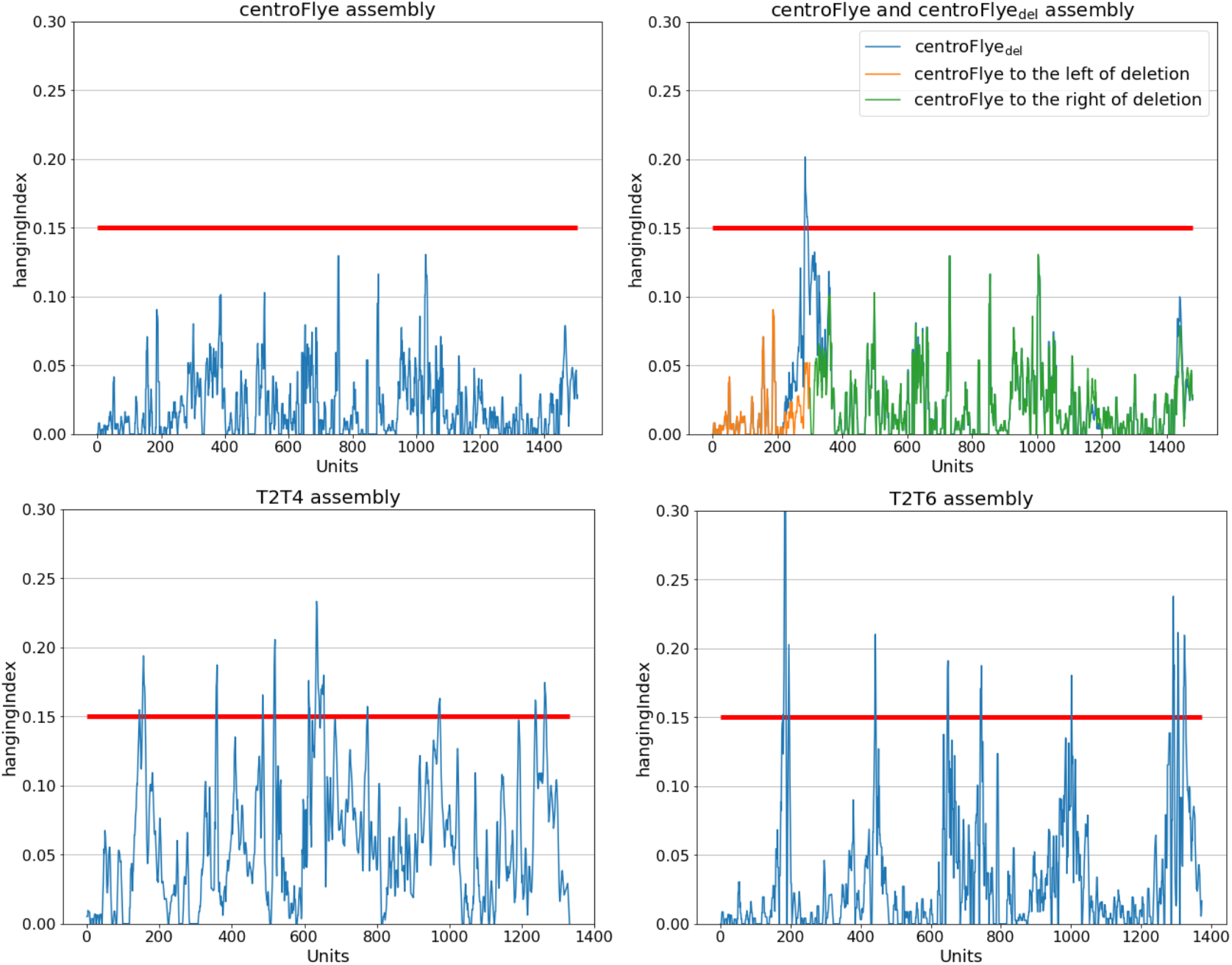
The hanging index test reveals putative breakpoints in centroFlye, centroFlye_del_, T2T4, and T2T6 assemblies. The peak at unit ~180 in T2T6 assembly **(bottom right)** is 0.4 and is cut. Red horizontal line at *y*=0.15 represents the default threshold for *hangingIndex*.

## Appendix K: Breakpoint test

Consider a read *Read* spanning positions from *a* to *b* in a centromere and a position *a* < *x* < *b.* We define *l(Read,x)* as the number of unique *k*-mers in region [*a,x*] of the centromere and *r(Read, x*) as the number of unique *k*-mers in [*x,b*] (we prefer this notation even though these numbers only depend on *a*, *x*, and *b* rather than *Read*). We further define *L(x)* (*R(x)*) as the sum of *l(Read,x) (r(Read,x))* over all reads *Read* that cover position *x*.

We also define *left(Read,x)* as the number of unique *k*-mers from [*a,x*] that appear in *Read* and *right(Read,x)* as the number of unique *k*-mers from [*x,b*] that appear in *Read.* We further define *Left(x) (Right(x))* as the sum of *left(Read,x) (right(Read,x))* over all reads *Read* that cover position *x.* Finally, we compute *leftRatio(x)*=*left(x)/L(x)* and *rightRatio(x)*=*right(x)/R(x)* for each position *x* in the reconstructed cenX and define *meanRatio(x)* as the average of *leftRatio(x) and rightRatio(x)*. We expect that *meanRatio* is approximately equal to *survivalRate* for all positions in cenX. Figure K1 shows the *meanRatio* functions for centroFlye, centroFlye_del_, T2T4, and T2T6 assemblies of cenX and reveals some drops in these assemblies.

Drops in values of *meanRatio* may point to breakpoints in cenX assembly caused by large multi-unit indels as illustrated by a drop for centroFlye_del_ assembly around the unit 300 (Figure K1, top right). All assemblies have a drop around units 650. For centroFlye assembly this is the only drop. The T2T4 assembly has significantly lower *meanRatio* (as compared to centroFlye and T2T6 assemblies), likely due to lack of polishing. T2T6 has five drops around units 180, 450, 650—750, 1000, and 1300. These drops are coordinated with the positions of mapping discrepancies between T2T6 and centroFlye assemblies (Figure 4), significantly inflated or deflated coverage in T2T6 assembly (5), positions of high discordance in T2T6 assembly (Figure 6), and positions with high *hangingIndex* in T2T assembly (Figure J1).

**Figure K1.**
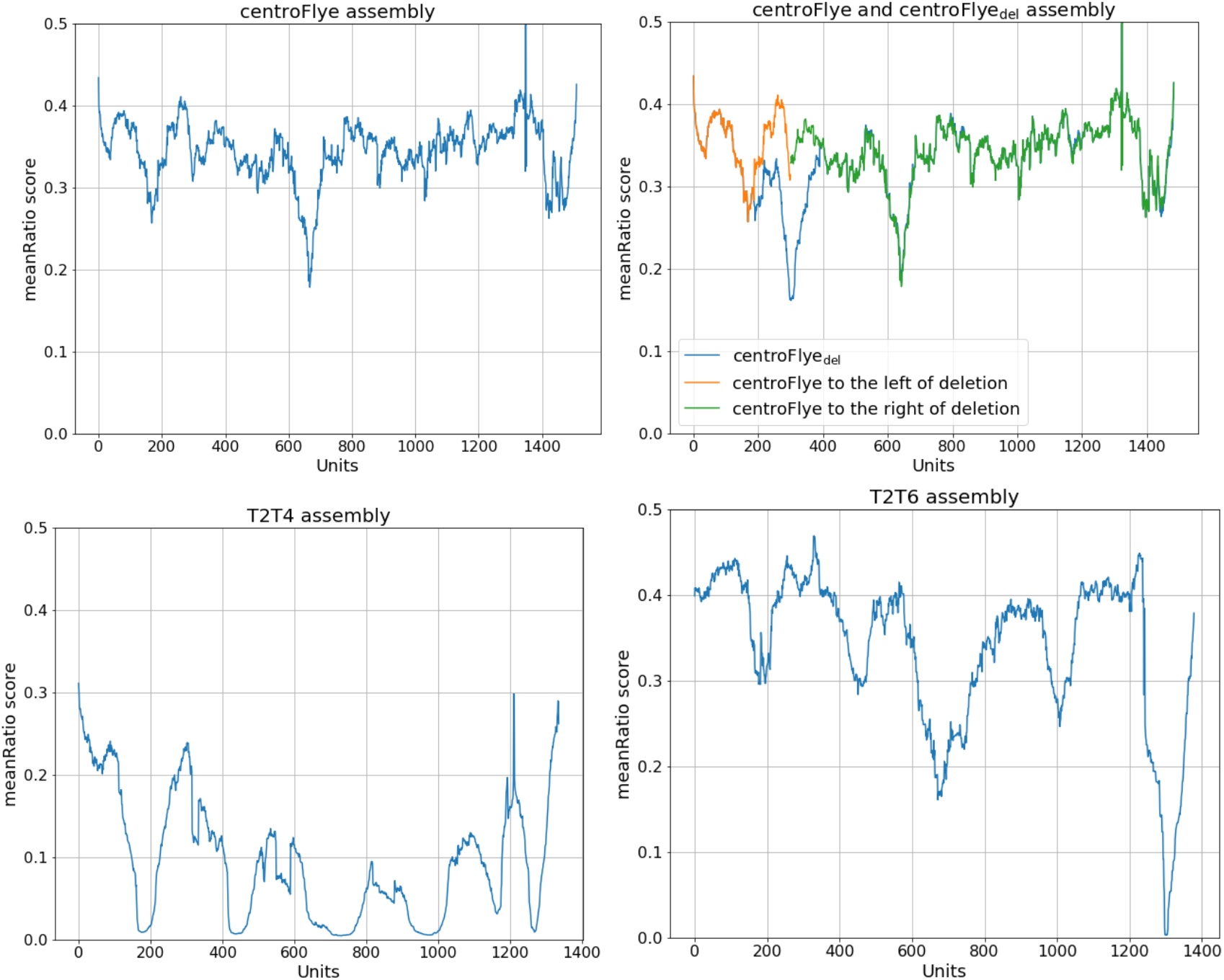
The breakpoint test reveals potential breakpoints in centroFlye, centroFlye_del_, T2T4, and T2T6 assemblies.

